# Signal peptide hydrophobicity modulates interaction with the twin-arginine translocase

**DOI:** 10.1101/135103

**Authors:** Qi Huang, Tracy Palmer

## Abstract

The general secretory pathway (Sec) and twin-arginine translocase (Tat) operate in parallel to export proteins across the cytoplasmic membrane of prokaryotes and the thylakoid membrane of plant chloroplasts. Substrates are targeted to their respective machineries by N-terminal signal peptides that share a common tripartite organization, however Tat signal peptides harbor a conserved and almost invariant arginine pair that are critical for efficient targeting to the Tat machinery. Tat signal peptides interact with a membrane-bound receptor complex comprised of TatB and TatC components, with TatC containing the twin-arginine recognition site. Here we isolated suppressors in the signal peptide of the Tat substrate, SufI, that restored Tat transport in the presence of inactivating substitutions in the TatC twin-arginine binding site. These suppressors increased signal peptide hydrophobicity, and co-purification experiments indicated that they restored binding to the variant TatBC complex. The hydrophobic suppressors could also act in *cis* to suppress substitutions at the signal peptide twin-arginine motif that normally prevent targeting to the Tat pathway. Highly hydrophobic variants of the SufI signal peptide containing four leucine substitutions retained the ability to interact with the Tat system. The hydrophobic signal peptides of two Sec substrates, DsbA and OmpA, containing twin lysine residues, were shown to mediate export by the Tat pathway and to co-purify with TatBC. These findings indicate that there is unprecedented overlap between Sec and Tat signal peptides and that neither the signal peptide twin-arginine motif nor the TatC twin-arginine recognition site are essential mechanistic features for operation of the Tat pathway.

**Importance:** Protein export is an essential process in all prokaryotes, The Sec and Tat export pathways operate in parallel, with the Sec machinery transporting unstructured precursors and the Tat pathway transporting folded proteins. Proteins are targeted to the Tat pathway by N-terminal signal peptides that contain an almost invariant twin-arginine motif. Here we make the surprising discovery that the twin-arginines are not essential for recognition of substrates by the Tat machinery, and that this requirement can be bypassed by increasing the signal peptide hydrophobicity. We further show that signal peptides of *bona fide* Sec substrates can also mediate transport by the Tat pathway. Our findings suggest that key features of the Tat targeting mechanism have evolved to prevent mis-targeting of substrates to the Sec pathway rather than being a critical requirement for function of the Tat pathway.

## Introduction

The general secretory (Sec) and twin-arginine translocation (Tat) pathways operate in parallel to transport proteins across the cytoplasmic membranes of prokaryotes and the thylakoid membranes of plant chloroplasts. The Sec pathway translocates substrates in an unfolded conformation, whereas the Tat system transports folded proteins. Many Tat substrates contain redox cofactors that are non-covalently associated, and the Tat system is essential for photosynthesis and some modes of respiratory growth (reviewed in (1)).

Targeting of substrates to the Sec and Tat pathways is mediated by the presence of N-terminal signal peptides. Sec and Tat targeting sequences each have a recognisable tripartite structure with a positively-charged n-region, a hydrophobic h-region and a polar c-region that usually contains a cleavage site for leader peptidase (2, 3; Fig 1A). One of the primary differences between them is the presence of an almost invariant arginine pair in the n-region of Tat signal peptides. These consecutive arginines are reported to be mechanistically essential for substrate translocation by the Tat pathway, and even conservative substitutions to lysine are poorly tolerated (e.g. 4, 5). Twin arginines, however, are also compatible with the Sec pathway and some Sec signal peptides have paired arginines in their n-regions. A second key difference is the relative hydrophobicity of the two types of signal peptide. Tat targeting sequences are notably less hydrophobic than Sec signal peptides, and increasing the hydrophobicity of the TorA signal peptide re-routes a passenger protein from Tat to Sec (6, 7). Finally one or more positive charges is frequently found in the c-region of Tat signals that is not mechanistically required for Tat transport but serves to block interaction of the signal peptide with the Sec pathway (6-9). None-the-less, despite these differences, over half of the *E. coli* Tat signal peptides that were tested showed some level of engagement with the Sec pathway when fused to a Sec-compatible reporter protein (10).

**Figure 1.**
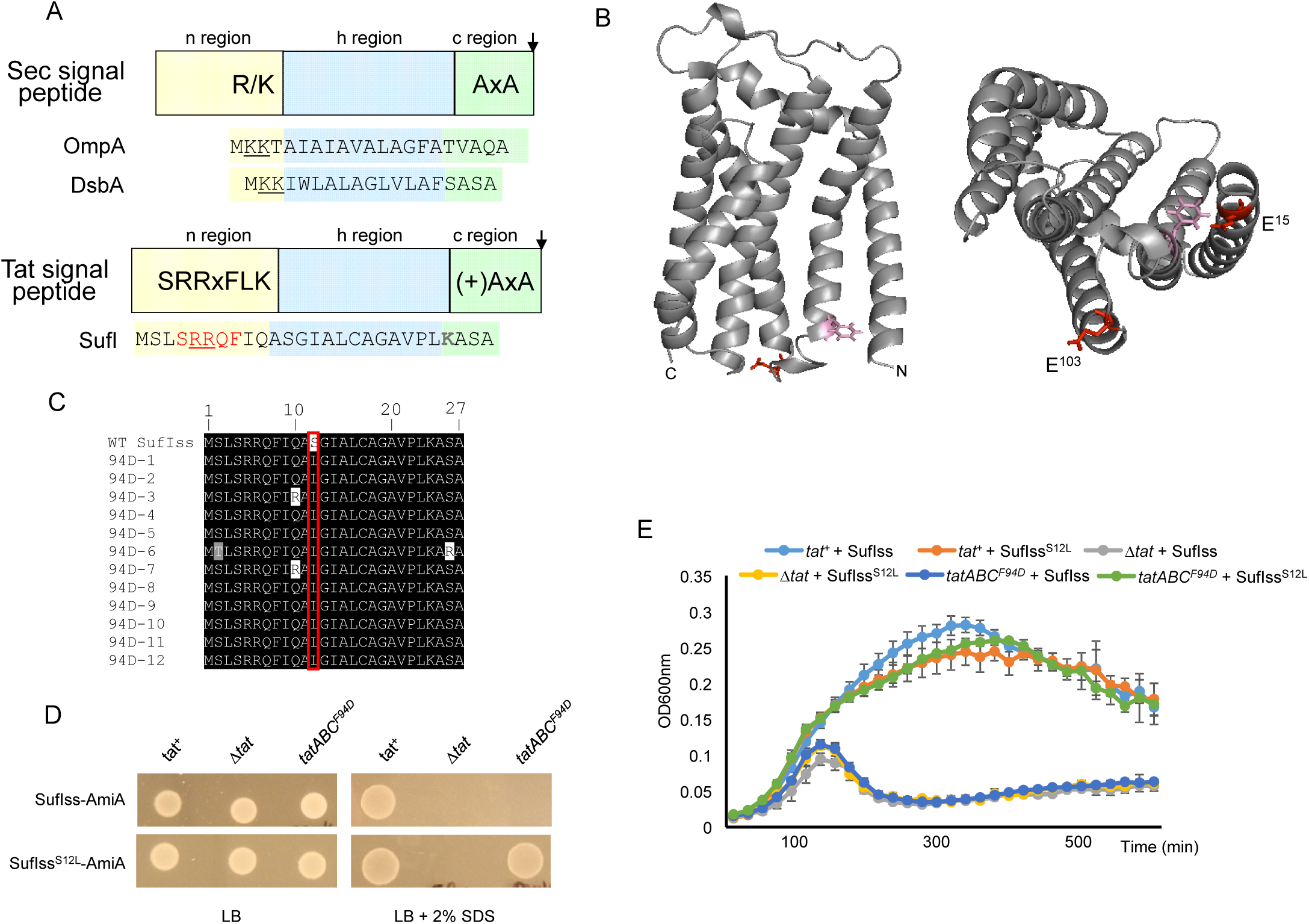
A. Schematic representation of Sec and Tat signal peptides. The sequence of the OmpA and DsbA Sec-targeting signals and the SufI Tat-targeting signal peptide are shown. Positive charges in the signal peptide n-regions are shown in underline, and the amino acids of the SufI Tat consensus motif are shown in red. B. Models of *E. coli* TatC (*left*) side view, with F^94^ and E^103^ residues that are located in the signal peptide binding site given in pink and red, respectively, and (*right*) view of the cytoplasmic face, with E^15^ additionally shown. C. Alignment of the amino acid sequence of twelve suppressors in the SufI signal peptide that compensate for the TatC F94D substitution. D and E. Cells of strain MC4100 Δ*amiA* Δ*amiC* Δ*tatABC* harboring pTH19kr (empty vector; annotated Δ*tat*) or pTAT101 producing wild type TatAB along with either wild type TatC (*tat*^+^) or TatC^F94D^ (*tatABC*^*F94D*^) and a compatible plasmid encoding either pSUSufIss-mAmiA or pSUSufI^S12L^ss-mAmiA, as indicated, were sub-cultured at 1:100 into fresh LB medium following overnight growth and: D. incubated for 3 hours at 37 °C with shaking. Cells were pelleted, re-suspended in sterile PBS supplemented with appropriated antibiotics to an OD_600_ of 0.1 and 8 μL of sample was spotted onto LB agar or LB agar containing 2% SDS Plates were incubated at 37 °C for 16 hours, or E. supplemented with 0.5% SDS (final concentration) and grown at 37 °C without shaking. The optical density at 600nm was monitored every 20min using a plate reader. Error bars are ± SD, *n*=3 biological replicates.

Tat signal peptides interact with the membrane-bound Tat receptor complex. In *E. coli* the receptor contains TatA, TatB and TatC proteins, most likely in a 1:1:1 stoichiometry (11-13). The receptor is multivalent (14-16) and contains multiple copies of the TatABC heterotrimer (e.g. 17, 18). The primary recognition site for the Tat signal peptide is TatC (e.g.19-23), with two conserved glutamates on the cytoplasmic face of TatC forming a patch that interacts with the signal peptide twin-arginines (24; Fig 1B). The signal peptide can also transition to a deep binding mode where it is inserted into the receptor complex, forming crosslinks to the transmembrane helix (TM) of TatB and TM5 of TatC (17, 19, 25, 26). Signal peptide insertion into the receptor drives structural reorganisation of the complex (13, 18, 26, 27) and the recruitment of further TatA molecules (19, 28-30). The assembled TatA oligomer mediates the transport of folded substrates across the membrane in an unknown manner, potentially by forming a translocation channel or by facilitating a localized weakening and transient disruption of the bilayer (31-33).

A recent study isolated genetic suppressors that restored transport activity to a Tat system that harbored an inactivating substitution in the TatC signal peptide binding site (27). These suppressing substitutions, located primarily in the TM of TatB, could also separately restore Tat transport to a substrate with a defective Tat signal peptide. Biochemical analysis revealed that these substitutions did not act to restore detectable signal peptide binding to the receptor complex but instead at least some of them induced conformational changes that apparently mimicked the substrate-activated state (27). In this work we have taken a complementary approach by searching for signal peptide suppressors able to restore Tat transport when the TatC signal peptide binding site was inactivated. We show that two separate inactive TatC variants, F94D and E103K, can be suppressed by single substitutions that increase the hydrophobicity of a Tat signal peptide. Remarkably, the same hydrophobic substitutions can suppress in *cis* by restoring Tat transport to a twin-arginine substituted signal peptide. Our results show that neither the twin-arginine motif nor its cognate recognition site on TatC are required for Tat transport activity. We further show that hydrophobic Sec signal peptides containing paired lysines can also mediate export by the Tat pathway pointing to an unexpected degree of overlap between Sec and Tat targeting requirements.

## Results

### Isolation of suppressors of the inactivating TatC F94D substitution

A series of crosslinking studies, along with direct binding assays using purified TatC variants, have identified that the cytoplasmic N-terminal region and the cytoplasmic loop between TM2 and TM3 forms a binding site for the twin-arginine motif of Tat signal peptides (23, 24, 34, 35). Amino acid substitutions in the TM2-TM3 loop in particular are associated with loss of Tat activity, and residues F94 and E103 are almost completely invariant among TatC sequences from all three domains of life (34, 36, 37). Along with E15, E103 has been implicated in co-ordinating the positively charged twin-arginines of the signal peptide (24, 38; Fig 1B).

The twin-arginines are part of a larger consensus motif, S-R-R-x-F-L-K (Fig 1A) where the other amino acids are semi-conserved (2). The consensus phenylalanine is frequently present, particularly in bacterial Tat signal peptides, and for example is found in approximately 2/3^rds^ of *E. coli* Tat targeting sequences (39). It has been proposed through modelling studies that if the signal peptide n-region is in an extended conformation, TatC^F94^ may stack against this consensus F residue (40). We initially sought to test this hypothesis genetically.

It has been shown previously that a TatC^F94D^ substitution inactivates Tat transport and that strains harboring this substitution are unable to grow on media containing the detergent SDS (27; Fig 1D). This phenotype arises due to an inability to export two Tat substrates, AmiA and AmiC that remodel the cell wall during growth (41, 42). We used a fusion protein whereby the signal peptide of SufI, which has the consensus F residue (Fig 1A) was fused to the mature region of AmiA (27) and constructed a random library of codon substitutions at F8. We then screened this library against a strain lacking native *amiA*/*amiC* and harboring TatC^F94D^, plating onto LB medium containing 2% SDS to select for suppressors of this inactivating substitution. However, after screening more than 1 000 clones we failed to isolate any suppressors of TatC^F94D^ from this library.

We therefore addressed whether it was possible to isolate substitutions elsewhere in the SufI signal peptide that would suppress TatC^F94D^. To this end we constructed a random library of mutations throughout the SufI signal peptide coding region of the SufIss-AmiA fusion that had some 13 000 members and an error rate of approximately 2%. After screening more than 20 000 individual transformants for the ability to grow in the presence of 2% SDS, we isolated 12 suppressors that supported growth on the detergent. Sequence analysis indicated that each of the suppressors shared a common substitution of serine at position 12 of the signal peptide to leucine (Fig 1C), and indeed this single S12L substitution was sufficient to support growth of a strain producing TatC^F94D^ on LB agar containing SDS (Fig 1D). Since the phenotypic growth test is largely qualitative, we also undertook a more quantitative assessment of growth of the strain co-producing TatC^F94D^ and SufI^S12L^ss-AmiA by measuring growth curves in the presence of SDS. Fig 1E shows that the strain producing TatC^F94D^ and SufI^S12L^ss-AmiA grew identically to the same strain producing wild type TatC and SufI^S12L^ss-AmiA.

### The SufI S12L substitution restore transport activity to a different substitution in the TatC signal peptide binding site

To determine whether the suppressor activity of the signal peptide S12L substitution was specific for TatC^F94D^ we tested whether this substitution could restore Tat transport to other TatC inactivating substitutions including P48L, V145E and Q215R located in consecutive periplasmic loops or E103K, located in the signal peptide binding site (37). Fig 2 shows that the inactivating TatC^E103K^ substitution could also be suppressed by the SufI^S12L^ variant, but transport activity was not restored to any of the substitutions in the periplasmic loops. We conclude that the S12L substitution specifically restores Tat transport to substitutions in the TatC signal peptide binding site.

**Figure 2.**
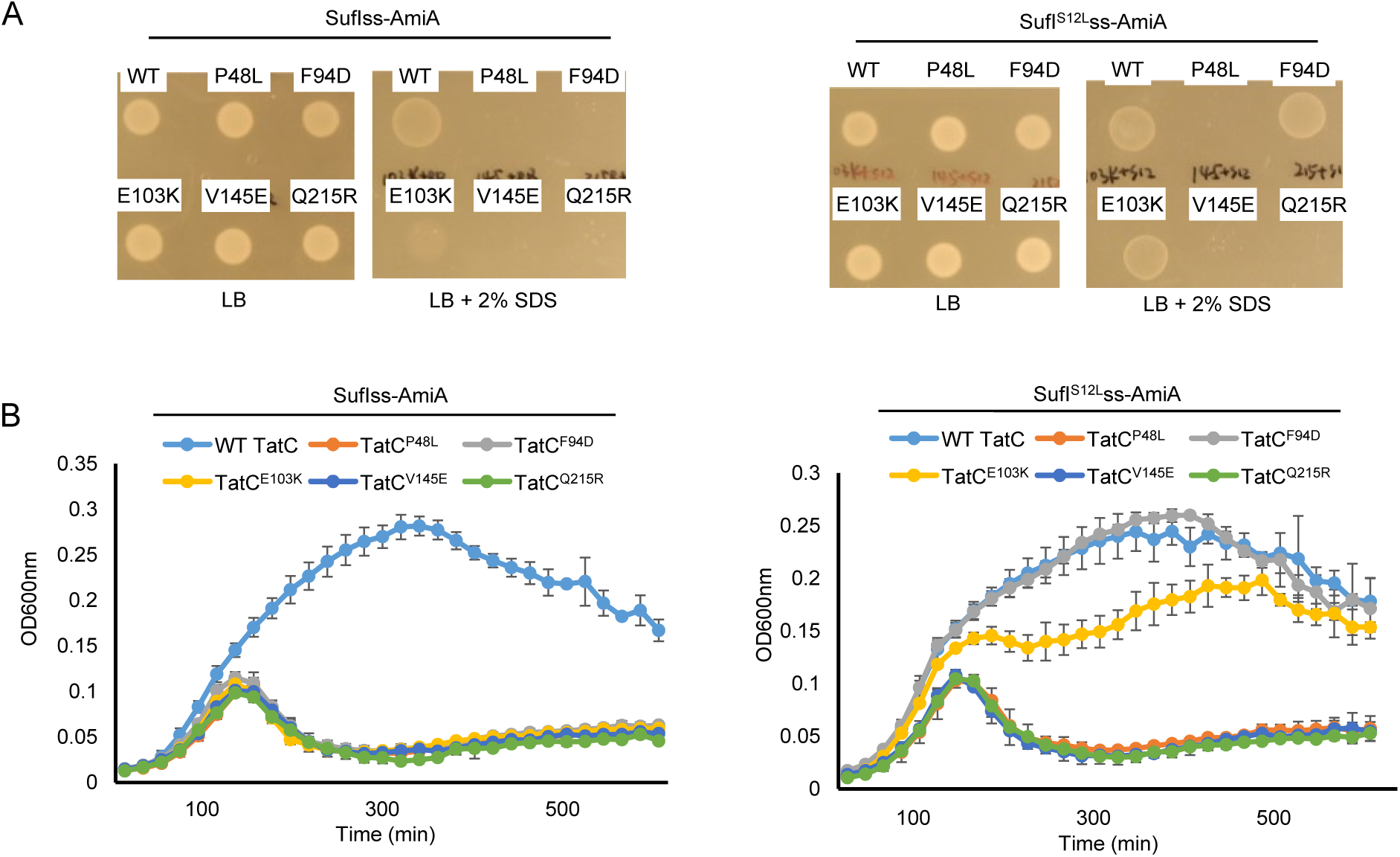
The SufI^S12L^ substitution can restore Tat transport to TatC^E103K^. A and B. Overnight cultures of strain MC4100 Δ*amiA* Δ*amiC* Δ*tatABC* harboring either pSUSufIss-mAmiA or pSUSufI^S12L^ss-mAmiA alongside plasmid pTAT101 producing wild type TatAB along with the indicated substitution of TatC, as indicated, were sub-cultured at 1:100 dilution and: A. grown for a further 3 hours at 37 °C, pelleted, re-suspended to an OD_600_ of 0.1 and 8 μL of sample was spotted onto LB agar or LB agar containing 2% SDS. Plates were incubated at 37 °C for 16 hours, or B. supplemented with 0.5% SDS (final concentration) and grown at 37 °C without shaking. The optical density at 600nm was monitored every 20min using a plate reader. Error bars are ± SD, *n*=3 biological replicates.

### The S12L substitution can restore transport activity to signal peptides that contain inactivating twin-arginine substitutions

Since the signal peptide S12L substitution can act in *trans* to suppress inactivating substitutions in the TatC signal peptide binding site, we next asked whether it could act in *cis* to rescue inactivating substitutions at the twin-arginine motif. Previously it has been shown that substitutions of one or both consensus arginines of the SufI signal peptide are poorly tolerated (4), and indeed single substitutions of R6 to D, E, H, N or Q, or of R5R6 to KK, KH, KQ or HH in the SufIss-AmiA fusion are sufficient to prevent phenotypic growth of cells in the presence of SDS (27; Fig 3, Fig S1). Interestingly, however, introduction of the S12L substitution alongside any of the R6D, R6E, R6H, R6N, R6Q, or R5K/R6K restored strong growth of cells producing these fusion proteins in the presence of SDS (Fig 3A, Fig S1 panels B-G). The S12L substitution could also partially compensate for the R5K,R6Q substitution (Fig 3A, Fig S1 panel H), but could not rescue transport activity of the R5K,R6H or R5H,R6H variants (Fig 3B, Fig S1, panels I and J). For each of these variant signal peptides we confirmed that transport of the AmiA substrate remained strictly Tat-dependent since growth on SDS was not observed when the Tat system was absent (Fig S2). We conclude that the SufI^S12L^ signal peptide substitution can at least partially compensate for substitutions at the twin-arginine motif.

**Figure 3.**
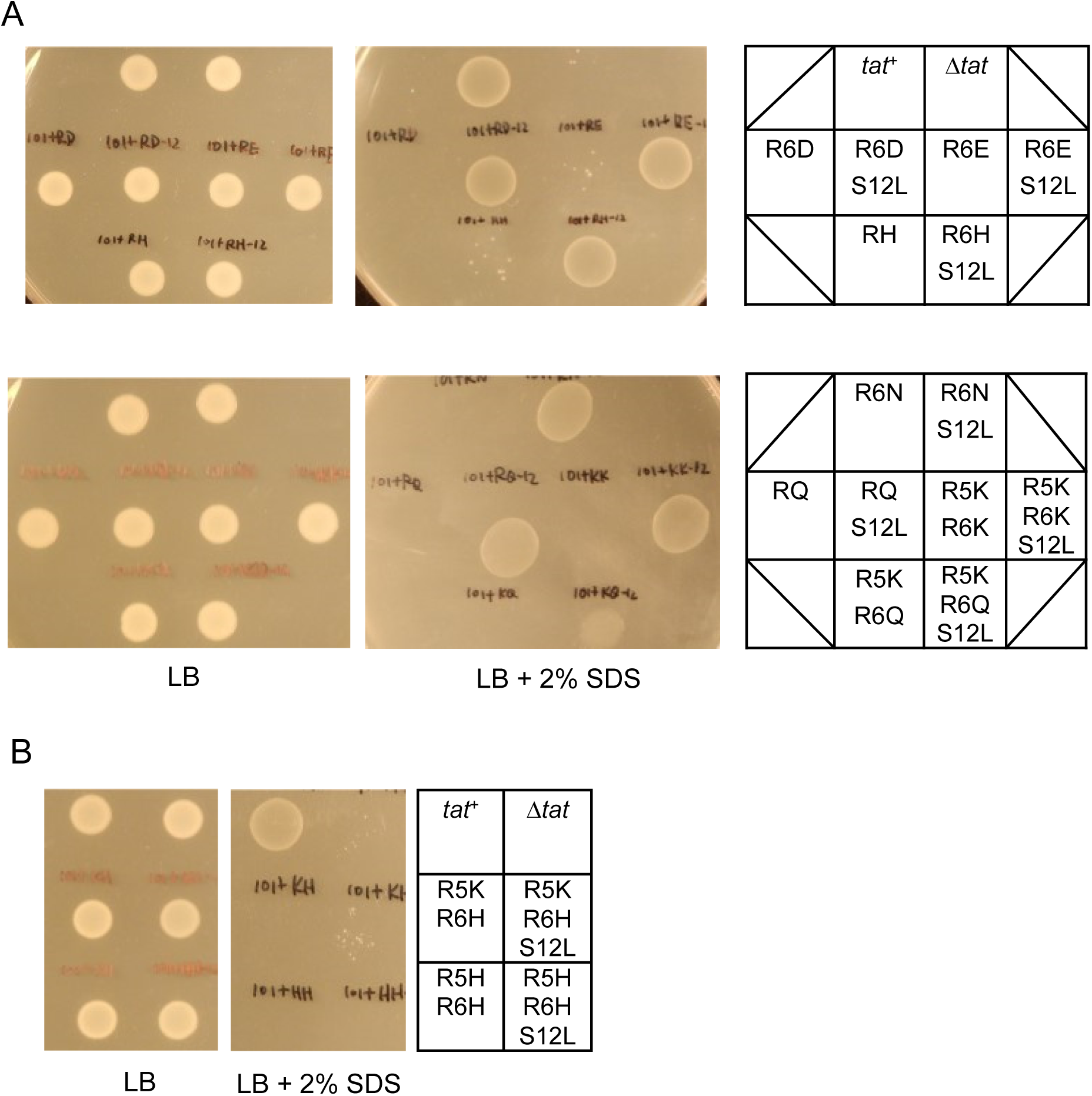
The SufI^S12L^ substitution can act in *cis* to suppress inactivating substitutions in the SufI signal peptide twin-arginine motif. A and B. Overnight cultures of strain MC4100 Δ*amiA* Δ*amiC* Δ*tatABC* harboring either pTH19kr alongside pSUSufIss-mAmiA (Δ*tat*) or pTAT101 (producing wild type TatABC) alongside either unsubstituted pSUSufIss-mAmiA (*tat*^+^) or pSUSufIss-mAmiA encoding the indicated substitutions in the SufI signal peptide were sub-cultured at 1:100 dilution and grown for a further 3 hours at 37 °C, pelleted, re-suspended to an OD_600_ of 0.1 and 8 μL of sample was spotted onto LB agar or LB agar containing 2% SDS. Plates were incubated at 37 °C for 16 hours.

### Single hydrophobic substitutions along the length the SufI signal peptide h-region can also suppress inactivating TatC substitutions in the signal peptide binding site

The h-regions of Tat signal peptides are less hydrophobic than Sec signal sequences, containing significantly more glycine and less leucine residues (6). The S12L substitution replaces a polar residue near the start of the SufI signal peptide h-region with a highly hydrophobic amino acid, markedly increasing its hydrophobicity score (Table 1). To test whether single hydrophobic substitutions elsewhere in the SufI signal peptide h-region could also suppress Tat transport defects, we increased hydrophobicity of the h-region by constructing individual A11L, G13L, A15L, A18L, G19L and A20L variants. Fig 4A shows that when each of these individual substitutions was introduced into the SufIss-AmiA construct and produced in a strain harboring *tatC*^*F94D*^, phenotypic growth on SDS was restored. When this was examined semi-quantitatively by following growth curves (Fig 4B), it could be seen that the G13L, A15L and G19L substitutions suppressed the *tatC* ^*F94D*^ allele more strongly than A11L, A18L or A20L. Table 1 shows that the substitutions that give the biggest increase in hydrophobicity result in the strongest level of suppression. It should be noted that the SufIss^G13L^ substitution appeared to result in a very low level of the fusion protein being routed to the Sec pathway as weak growth could be detected in a strain lacking the Tat pathway (Fig 4A, Fig 4C). None of the other substitutions, however, led to any detectable transport by Sec.

**Figure 4.**
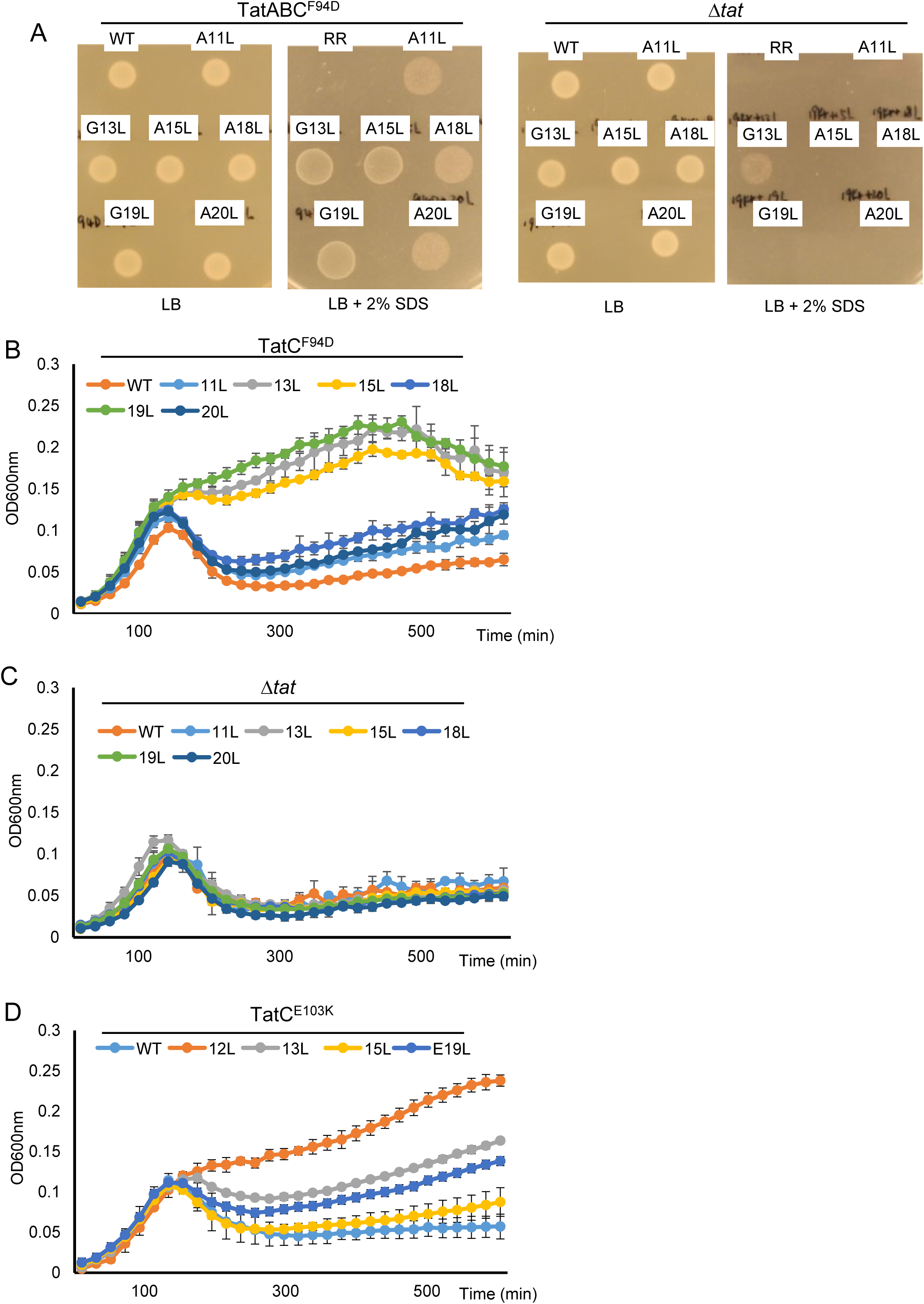
Single leucine substitutions throughout the SufI signal peptide h-region can suppress the TatC F94D and E103K substitutions. A - C. Overnight cultures of strain MC4100 Δ*amiA* Δ*amiC* Δ*tatABC* harboring either pTH19kr (Δ*tat*) or pTAT101 producing TatABC^F94D^ along with pSUSufIss-mAmiA producing the indicated substitution in the SufI signal peptide were sub-cultured at 1:100 dilution and: A. grown for a further 3 hours at 37 °C, pelleted, re-suspended to an OD_600_ of 0.1 and 8 μL of sample was spotted onto LB agar or LB agar containing 2% SDS. Plates were incubated at 37 °C for 16 hours, or B. and C. supplemented with 0.5% SDS (final concentration) and grown at 37 °C without shaking. D. MC4100 Δ*amiA* Δ*amiC* Δ*tatABC* harboring pTAT101 producing TatABC^E103K^ along with pSUSufIss-mAmiA producing the indicated substitution in the SufI signal peptide was subcultured into LB containing 0.5% SDS and grown at 37 °C without shaking. For all growth curves the optical density at 600nm was monitored every 20min using a plate reader. Error bars are ± SD, *n*=3 biological replicates.

**Table 1.** Relative hydrophobicities of signal peptide variants used in this work. In each case the h-region sequence used to calculate the score is shown underlined. Hydrophobicity was scored using grand average of hydropathy (GRAVY) value at http://www.gravy-calculator.de/.

| Signal peptide | Sequence | Hydrophobicity |
| --- | --- | --- |
| WT SufI | MSLSRRQFIQ<u>ASGIALCAGAVPL</u>KASA | 1.75 |
| SufI 11L | MSLSRRQFIQ<u>L</u>SGIALCAGAVPLKASA | 1.91 |
| SufI 12L | MSLSRRQFIQ<u>AL</u>GIALCAGAVPLKASA | 2.11 |
| SufI 13L | MSLSRRQFIQ<u>ASL</u>IALCAGAVPLKASA | 2.08 |
| SufI 15L | MSLSRRQFIQ<u>ASGILL</u>CAGAVPLKASA | 1.92 |
| SufI 18L | MSLSRRQFIQ<u>ASGIALCL</u>GAVPLKASA | 1.91 |
| SufI 19L | MSLSRRQFIQ<u>ASGIALCAL</u>AVPLKASA | 2.08 |
| SufI 20L | MSLSRRQFIQ<u>ASGIALCAGLV</u>PLKASA | 1.91 |
| SufI 12L,13L | MSLSRRQFIQ<u>ALL</u>IALCAGAVPLKASA | 2.43 |
| SufI 12L,13L,14L,15L | MSLSRRQFIQ<u>ALLLLL</u>CAGAVPLKASA | 2.53 |
| SufI 17L,18L,19L,20L | MSLSRRQFIQ<u>ASGIALLLLL</u>VPLKASA | 2.48 |
| WT OmpA | MKKTAIAIAVALAGFATVAQA | 2.52 |
| OmpA i18K | MKKTAIAIAVALAGFATKVAQA | 2.52 |
| WT DsbA | MKKIWLALAGLVLAFSASA | 2.57 |
| DsbA i16K | MKKIWLALAGLVLAFKSASA | 2.57 |

We next tested whether these further hydrophobic substitutions could also suppress a second signal peptide binding site substitution, TatC^E103K^. It can be seen (Fig S3A, Fig 4D) that these variants could also compensate for loss of Tat activity resulting from this substitution. They could not, however, compensate for any of TatC P48L, V145E and Q215R (not shown). As seen for the suppression of TatC^F94D^, the substitutions giving the biggest increase in hydrophobicity (S12L, G13L and G19L) restored the highest level of Tat activity in the presence of TatC^E103K^ (Fig 4D). Finally, we also tested whether the strongest suppressors of TatC^E103K^ could also rescue transport in *cis* when introduced into the twin lysine variant of the SufI signal peptide. Fig S3B indicates that similar to the S12L substitution, introduction of any of the G13L, A15L and G19L substitutions into KK-SufIss-AmiA restored strong growth of cells producing these fusion proteins in the presence of SDS. We conclude that increasing h-region hydrophobicity can suppress transport defects associated with either the signal peptide twin-arginine motif or the signal peptide binding site.

### The h-region suppressors support transport of full-length SufI

To assess the level of Tat transport mediated by these hydrophobic variants of the SufI signal sequence, we introduced the S12L, G13L, A15L and G19L substitutions individually into a construct encoding C-terminally His-tagged but otherwise wild-type SufI. We initially expressed these in a strain producing native TatABC and fractionated cells to obtain the periplasm. Fig 5A shows that each of these single hydrophobic variants of the SufI signal peptide supported strong export of SufI-His. Transport of these SufI-His variants was strictly dependent on the Tat pathway since no periplasmic SufI-His could be detected when the Tat pathway was absent (Fig S4A).

**Figure 5.**
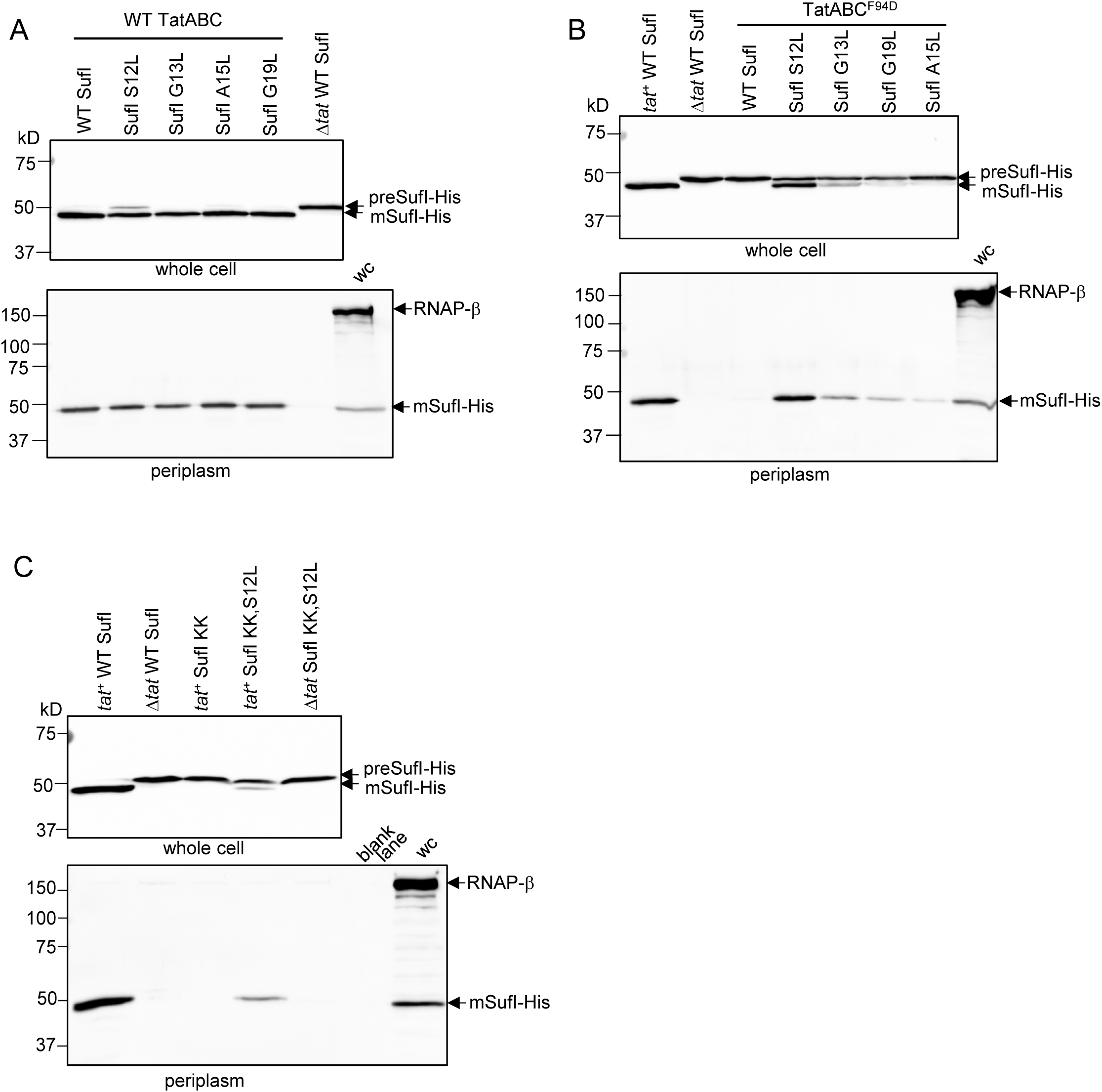
Analysis of SufI export mediated by signal peptide leucine substitutions in the TatC^F94D^ background or when combined with a signal peptide twin-lysine substitution. A. and *E. coli* strain DADE co-producing his-tagged but otherwise native SufI, or SufI with the indicated single leucine substitutions in the signal peptide (from a pQE80 plasmid) alongside either A. wild-type TatABC or B. wild type TatAB and TatC^F94D^ (from pTAT101). Strain DADE co-producing his-tagged but otherwise native SufI alongside an empty vector was used as a negative control (lanes annotated ‘Δ*tat* WT SufI’), or C. Strain DADE producing his-tagged SufI harboring an R5K,R6K double substitution (SufI KK) and with an additional S12L substitution where indicated, alongside either empty vector pTH19kr (Δ*tat*) or pTAT101 encoding wild type TatABC (*tat*^+^). In each case strains were grown to mid-log phase and fractionated into whole cell (upper panels) and periplasm (lower panels), then analyzed by Western blot with anti-6X His tag® or anti-RNA polymerase β subunit antibodies (cytoplasmic control protein). wc – whole cell. Equivalent volumes of sample were loaded for each of the whole cell samples, and for each of the periplasmic samples.

Next we assessed the degree of transport mediated by these variant signal peptides in cells producing TatABC^F94D^. It can be seen (Fig 5B) that although wild type SufI-His was not exported in the presence of TatC^F94D^, transport was detected when any of the single hydrophobic substitutions were present in the signal peptide. The S12L substitution in particular could strongly suppress TatC^F94D^, with high levels of SufI-His detected in the periplasm when the signal peptide harbored this mutation. These same signal peptide substitutions could also restore good transport of SufI-His in the presence of the inactivating TatC^E103K^ substitution (Fig S4B).

Since the hydrophobic substitutions can act in *cis* to restore transport activity to a twin lysine variant of the SufI signal peptide twin-arginine motif, we assessed the export of the KK variant of SufI-His harboring the S12L substitution. Fig 5C shows that there was clear Tat transport activity conferred on the twin lysine signal peptide variant by the presence of the S12L suppressor. Taken together the results presented so far indicate that the signal peptide consecutive arginines and the TatC twin-arginine recognition site are not essential mechanistic features for operation of the Tat pathway and substitutions in either of these can be at least partially compensated for by an increase in signal peptide hydrophobicity.

### The h-region suppressors restore signal peptide binding to TatBC

A previous study identified suppressors in the TatB component that could also restore transport activity to substitutions in the TatC twin-arginine binding site. It was shown that substrate precursors could be co-purified with wild type TatBC complexes but did not co-purify when the signal peptide binding site was mutated, even in the presence of the TatB suppressors. Thus it was concluded that the TatB suppressors did not detectably restore binding of signal peptides to the TatBC complex (27). To determine whether the suppressors we identified here that increase signal peptide hydrophobicity could restore binding to TatBC complexes harboring the TatC^F94D^ substitution, we co-produced FLAG-tagged variants of SufI with these suppressors alongside TatB and His-tagged TatC. After purification of TatBC complexes from digitonin-treated cell lysates, we assessed the level of co-purifying SufI by immunoblotting. As shown in Fig 6, the single substitutions S12L, G13L, A15L or G19L in the SufI signal peptide did not detectably affect interaction of SufI with wild type TatBC, since qualitatively similar levels of FLAG-tagged SufI were seen to co-purify with TatBC-His. When the F94D substitution was present in His-tagged TatC, no SufI-FLAG was co-purified with the variant TatBC-His complex (Fig 6A), even though SufI was clearly detected in the cell lysate (Fig S5). However, when the S12L, G13L, A15L or G19L substitutions were introduced into SufI, it could now be detected in the fractions containing purified TatBC^F94D^-His. These observations indicate that the SufI h-region suppressors restore some degree of signal peptide binding to the TatBC^F94D^ complex.

**Figure 6.**
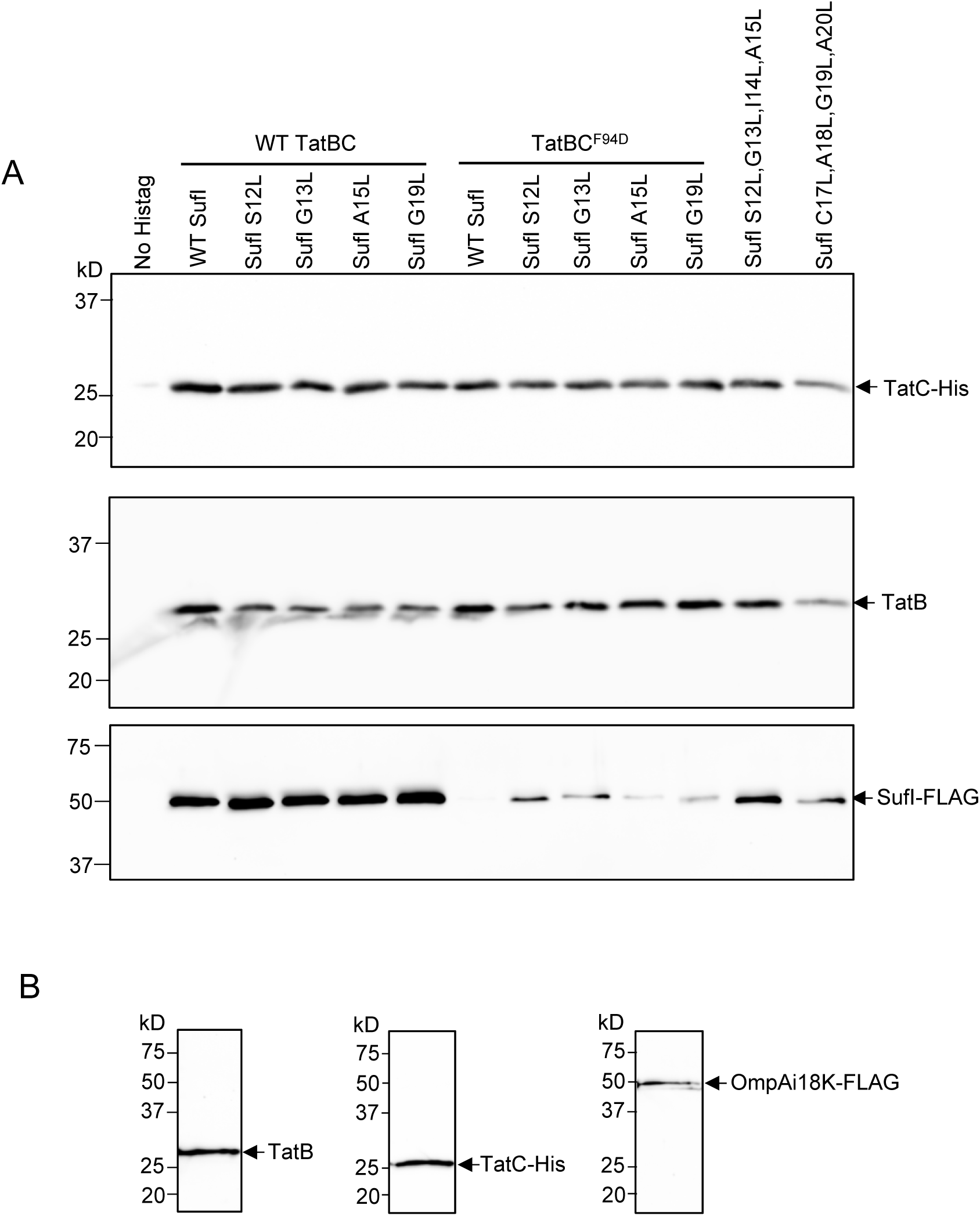
Hydrophobic variants of the SufI signal peptide mediate binding to TatBC and TatBC^F94D^. A. Cells of strain DADE-P co-producing C-terminally FLAG-tagged SufI with its native signal peptide (WT SufI) or harboring the indicated leucine substitutions, alongside TatB and C-terminally His-tagged TatC or TatC^F94D^ were lysed and incubated with digitonin and His-tagged TatC was isolated with Ni-charged beads. Following elution of bound TatC-His, equivalent volumes of the eluate from each sample were analyzed by western blotting with anti-His, anti-TatB, or anti-FLAG antibodies as indicated. DADE-P co-producing C-terminally FLAG-tagged SufI with its native signal peptide alongside TatB and non-tagged TatC (lane annotated ‘No Histag) was used as a negative control. Equivalent volumes of sample were loaded in each lane. B. Cells of DADE-P co-producing C-terminally FLAG-tagged SufI fused to the OmpA signal peptide harboring a lysine insertion at codon 18, alongside TatB and C-terminally His-tagged TatC were treated as described in A and equivalent volumes of the elution fraction were analyzed by western blotting with anti-His, anti-TatB, or anti-FLAG antibodies as indicated.

### Highly hydrophobic signal peptides are compatible with the Tat pathway

It has previously been reported that the relatively low hydrophobicity of Tat signal peptides partially prevents their routing to the Sec pathway (6). It is not clear, however, whether low h-region hydrophobicity is a mechanistic requirement for engagement with Tat. To explore this in more detail, we investigated the effect of further increasing the hydrophobicity of the SufI signal peptide on transport of the SufIss-AmiA fusion. To this end we introduced a S12L/G13L double substitution, and two quadruple substitutions, S12L/G13L/I14L/A15L and C17L/A18LG19L/A20L, into the SufI signal sequence. These substitutions markedly increase the signal sequence hydrophobicity score, bringing it into the range of the Sec signal sequences of OmpA and DsbA (Table 1).

Fig 7A shows that each of these SufIss-AmiA fusion proteins was able to support growth on solid medium in the presence of SDS, although growth was also seen in a strain lacking the Tat pathway, indicating that there is some export of these more hydrophobic SufIss-AmiA fusion proteins by Sec. These findings were confirmed by monitoring growth of these strains in liquid culture (Fig 7B,C). However it is clear that in the absence of a functional Tat system, growth in the presence of SDS was much poorer than when the Tat system was present. This observation suggests that there must be some recognition of these hydrophobic signal peptide variants by the Tat pathway. To confirm this, we co-produced FLAG-tagged S12L/G13L/I14L/A15L and C17L/A18LG19L/A20L SufI variants alongside TatBC-His. When His-tagged TatC was purified from digitonin-solubilized cell lysates, each of these hydrophobic SufI-FLAG variants was co-purified (Fig 6A), indicating that they retained the ability to interact with the TatBC complex. Taken together, these results show that highly hydrophobic signal peptides are mechanistically compatible with the Tat pathway.

**Figure 7.**
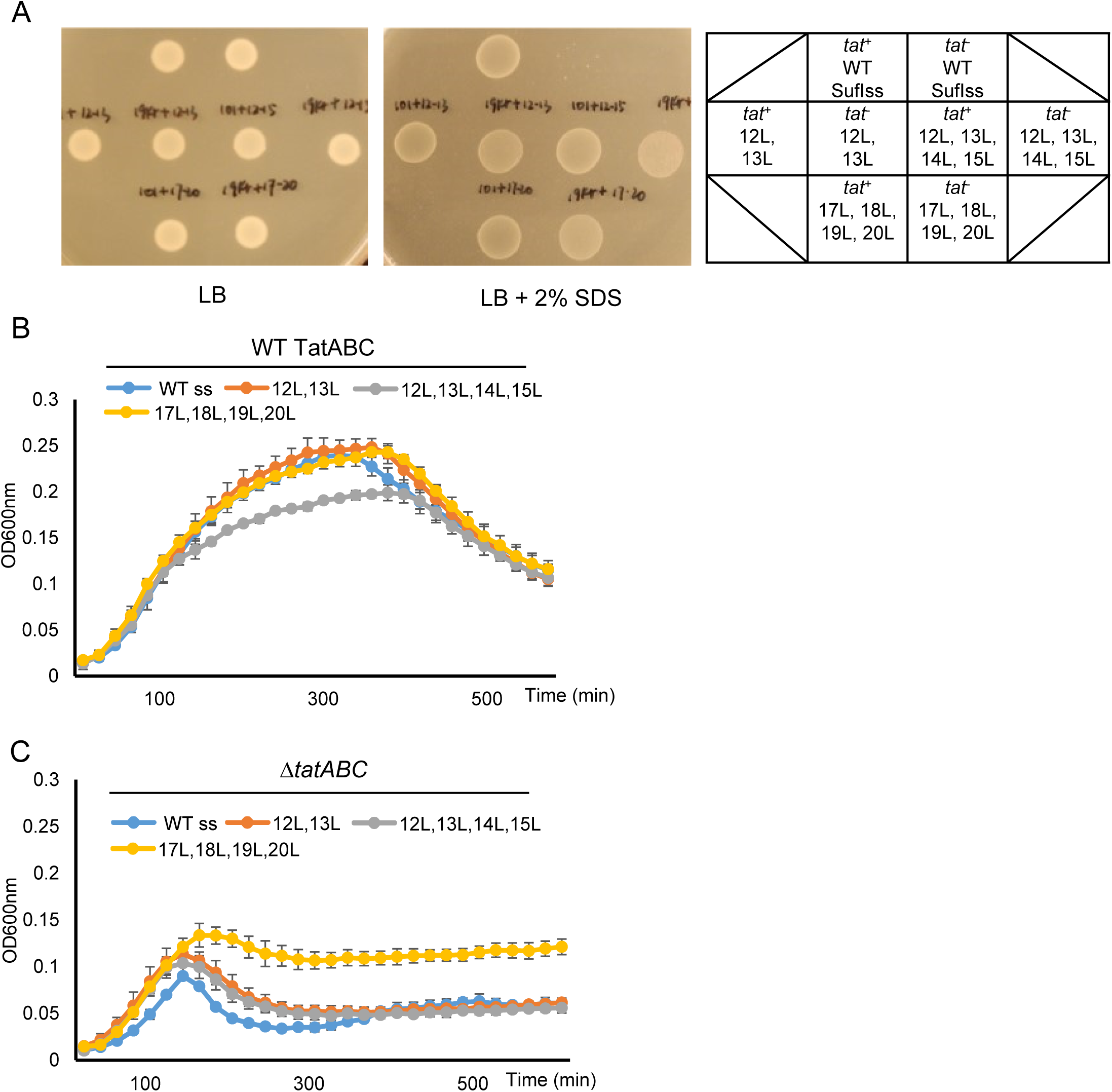
Multiple leucine substitutions in SufI signal peptide h-region partially re-route AmiA to the Sec pathway. A - C. Overnight cultures of strain MC4100 Δ*amiA* Δ*amiC* Δ*tatABC* harboring either pTH19kr (Δ*tat*) or pTAT101 producing wild type TatABC (*tat*^+^) along with pSUSufIss-mAmiA producing the indicated substitutions in the SufI signal peptide were sub-cultured at 1:100 dilution and: A. grown for a further 3 hours at 37 °C, pelleted, re-suspended to an OD_600_ of 0.1 and 8 μL of sample was spotted onto LB agar or LB agar containing 2% SDS. Plates were incubated at 37 °C for 16 hours, or B. and C. supplemented with 0.5% SDS (final concentration) and grown at 37 °C without shaking. The optical density at 600nm was monitored every 20min using a plate reader. Error bars are ± SD, *n*=3 biological replicates.

### The OmpA and DsbA signal peptides functionally interact with the Tat pathway

Our results collectively show that the hallmark twin-arginines of Tat signal peptides are not a mechanistic requirement for Tat-dependent transport and that a single arginine or twin lysines in the n-region are compatible with the Tat pathway if compensatory mutations are introduced that increase the hydrophobicity of the signal peptide. Interestingly, many Sec-dependent signal peptides share these parameters (Table S1), raising the possibility that *bona fide* Sec signal peptides may be able to interact with the Tat pathway. To explore this we selected two well-studied Sec signal peptides – those of OmpA, which is a post-translational Sec substrate and of DsbA which directs co-translational translocation (43, 44; Table 1) - and fused their signal peptides to the mature portion of AmiA. We also made two additional constructs where we introduced a ‘Sec-avoidance’ lysine residue into the signal peptide c-regions, to reduce interaction with the Sec pathway (8).

Fig 8A shows that there is Sec-dependent transport of AmiA mediated by the OmpA signal peptide as there is strong growth of the Δ*tat* strain producing OmpAss-AmiA in the presence of SDS. Introduction of a lysine at position 18 of the OmpA signal peptide clearly reduces interaction with the Sec pathway as growth of the Δ*tat* strain producing this variant is significantly reduced. However, there is good growth of *tat*^+^ strain producing this variant fusion protein, indicating that some of this fusion must be interacting with the Tat pathway. Similarly, Fig 8B shows that there is some low level growth of the Δ*tat* strain producing DsbAss-AmiA in SDS-containing medium, which is reduced by inclusion of a Sec-avoidance lysine in the c-region of the DsbA signal peptide. By contrast, the *tat*^+^ strain harboring either of these fusion proteins shows markedly stronger growth in the presence of SDS, indicating that the DsbA signal peptide is productively engaging with the Tat machinery.

**Figure 8.**
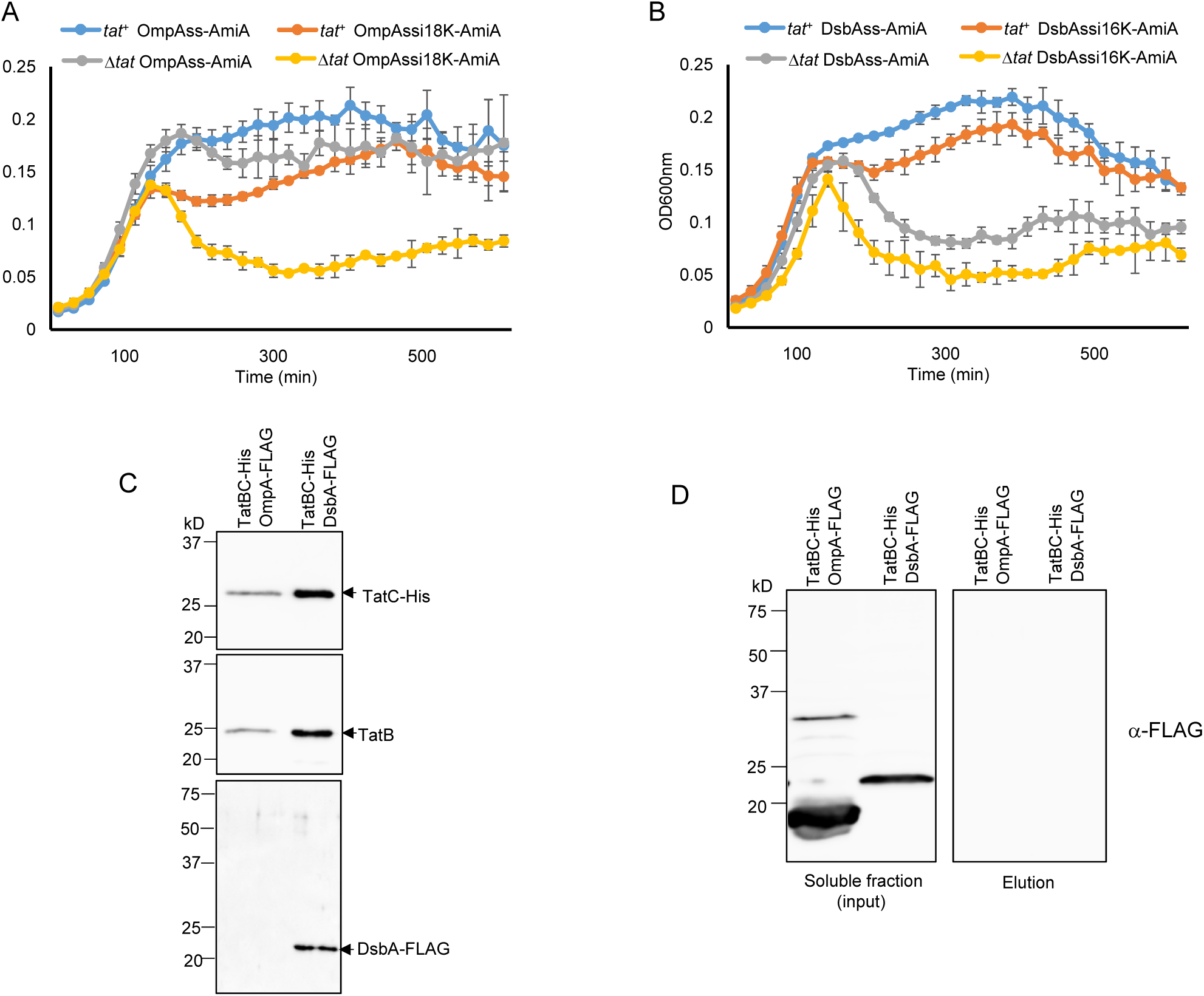
The OmpA and DsbA signal peptides are able to functionally engage with the Tat machinery. A. and B. Overnight cultures of strain MC4100 Δ*amiA* Δ*amiC* Δ*tatABC* harboring either pTH19kr (Δ*tat*) or pTAT101 producing wild type TatABC (*tat*^+^) along with a plasmid encoding the indicated signal peptide fusion to AmiA were sub-cultured at 1:100 dilution, supplemented with 0.5% SDS (final concentration) and grown at 37 °C without shaking. The optical density at 600nm was monitored every 20min using a plate reader. Error bars are ± SD, *n*=3 biological replicates. C. Cells of strain DADE co-producing TatB, C-terminally His-tagged TatC and either C-terminally FLAG-tagged DsbA or OmpA, as indicated, were lysed and incubated with digitonin and His-tagged TatC was isolated using Ni-charged beads. Following elution of bound TatC-His, equivalent volumes of the eluate from each sample were analyzed by western blotting with anti-His, anti-TatB, or anti-FLAG antibodies. D. An aliquot cell lysate prior to digitonin treatment from part C. was ultracentrifuged to remove the cell membranes. A small amount of the amount of the supernatant was retained as the input fraction and the remainder was incubated with Ni-charged beads. The beads were washed three times with wash buffer and aliquots of the input and eluate samples were analyzed by western blotting using an anti-FLAG antibody.

To confirm that these signal peptides are able to interact with Tat, we co-produced C-terminally FLAG-tagged variants of full length OmpA or full length DsbA alongside TatBC-His. When His-tagged TatC was purified from digitonin-solubilized cell lysates, DsbA-FLAG, which migrated very close to the expected mass of 24.1KDa, was seen to co-purify (Fig 8C). This co-purification was clearly dependent on the presence of the Tat proteins since when membranes were removed by an ultra-centrifugation step, the cytoplasmic form of DsbA-FLAG was no longer isolated by Ni-affinity purification (Fig 8D). We conclude that FLAG-tagged, but otherwise native DsbA can interact with TatBC. By contrast we were not able to detect co-purification of FLAG-tagged OmpA with TatBC under these conditions (Fig 8C). However in these experiments we noted that OmpA-FLAG migrated at a lower mass than the predicted size of the tagged protein (38.2kDa), or of folded OmpA (which migrates at an estimated mass of approximately 30kDa; (45)), raising the possibility that it may have been subjected to proteolysis. We therefore took a second approach to assessing whether the OmpA signal peptide could interact with TatBC by fusing the OmpA signal peptide variant containing the K18 insertion to the N-terminus of mature SufI, and co-producing it with TatBC-His. Fig 6C indicates that this fusion protein could indeed be co-purified alongside TatB and His-tagged TatC indicating that the OmpA signal peptide is able to interact with the TatBC complex.

## Discussion

In this study we have sought to identify SufI signal sequence variants that restore Tat transport activity in the presence of substitutions that inactivate the twin-arginine recognition site on TatC. Our results have shown that an increase in signal peptide hydrophobicity can overcome two different inactivating substitutions, TatC^F94D^ and TatC^E103K^, and that these suppressors act to restore detectable binding of the SufI signal sequence to the variant TatBC^F94D^ complex. We further showed that the same hydrophobic substitutions can act in *cis* to compensate for a range of inactivating substitutions at the SufI signal peptide twin-arginine motif. These results demonstrate that neither the consecutive arginines of the signal peptide nor the conserved recognition site on the cytoplasmic surface of TatC are mechanistically essential for operation of the Tat pathway, and that they can be bypassed if the signal peptide hydrophobicity is increased. Taken together our findings indicate that the signal peptide features that can facilitate interaction with the Tat pathway are remarkably similar to those that facilitate interaction with Sec, namely the presence of at least one basic charge in the n-region, and a relatively hydrophobic h-region. Indeed we show that even a highly hydrophobic signal peptide that naturally directs its passenger into the co-translocational Sec pathway can functionally engage with the Tat system.

If the Tat pathway can interact with hydrophobic signal peptides lacking the twin-arginine motif, why then do almost all Tat substrates that have been identified contain paired arginine residues and only moderately hydrophobic h-regions? In prokaryotes and plant chloroplasts, the Tat system always co-exists with the Sec pathway. In bacteria, ribosomal-associated signal recognition particle (SRP) and cytosolic or ribosomal-bound SecA capture Sec substrates at an early stage of biogenesis, at least partially through interaction with their signal sequences (46, 47). Signal sequence hydrophobicity is key sorting feature for Sec substrates; highly hydrophobic signals generally interact with SRP whereas those with lower hydrophobicity bind to SecA (46). Photocrosslinking and/or genetic studies have indicated that Tat signal peptides interact with ribosomally-bound trigger factor and with general cytoplasmic chaperones including DnaK (48-51), but no crosslinks to SecA have been reported, and *in vitro* analysis indicates that Tat signal peptides do not productively engage with SecA to the same extent as a Sec signal peptide (52). It is therefore likely that Tat signal peptides evolved lower hydrophobicity to avoid the targeting pathways that feed into the Sec translocon, and that the paired arginines and the twin-arginine binding site are necessary features to strengthen recognition of these weakly hydrophobic signal peptides by the Tat machinery. In this context it is interesting to note that although paired arginines in the signal peptide n-region are compatible with the Sec pathway, this pairing is relatively rare in Sec signal peptides, at least in *E. coli*, being found in only five of the 244 probable Sec signal peptides listed in Table S1 (compared with 53 that have paired lysines). If lysine and arginine are equivalent in the amino terminal region of a Sec signal peptide, as implied by kinetic analysis (53), this might suggest that there is selection pressure against the presence of paired arginines in Sec signals.

The presence of one or more positively charged amino acids in the c-region of Tat signal peptides is a further feature that has no mechanistic requirement for Tat translocation but leads to rejection of these signal sequences by the Sec pathway (6, 8, 9). C-terminal positive charges may act at a late stage during Sec translocation when the signal peptide is already engaged with the Sec translocon, and Sec avoidance motifs are particularly abundant in membrane proteins that require the dual action of the Sec and Tat pathways for their assembly (7, 54). Here the Tat-dependent signal sequence (which is internal to the protein and follows a series of Sec-dependent transmembrane domains) has several c-region positive charges that result in abortive interaction with the Sec pathway, freeing up the sequence to be recognized by Tat (7). Taken together it is clear that there is strong selective pressure, particularly at the level of Tat signal peptides, to refine features that minimise mis-targeting to the Sec pathway.

Our findings show that signal peptides with either twin lysines or an unpaired arginine, coupled with a moderately hydrophobic h-region can functionally interact with the Tat pathway. Inspection of all of the signal peptides present at the N-termini of *E. coli* MG1655 proteins identified using SignalP 4.1 (http://www.cbs.dtu.dk/services/SignalP; 55) indicates that some 44% of Sec signal peptides contain either KR, RK, KK, RD, RE, RH, RN, RQ adjacent to their h-regions (Table S1) and therefore potentially have the capability of engaging with the Tat pathway. Whether any of these would ever target to the Tat pathway *in vivo* is not clear, since presumably under standard conditions *E. coli* synthesizes sufficient targeting factors to ensure that Sec substrates are efficiently channelled into the Sec pathway. However, it should be noted that in *Bacillus subtilis*, hyper-production of a normally Sec-dependent lipase results in overflow into the Tat pathway (56), raising the possibility that transient re-routing of substrates to the Tat pathway may occur on occasions where cells undergo secretion stress.

## Materials and Methods

### Strain and plasmid construction

Strains used in this study are MC4100 derivatives (57). Strain MC4100 Δ*amiA* Δ*amiC* Δ*tatABC* (F-Δ*lacU169 araD139 rpsL150 relA1 ptsF rbs flbB5301* Δ*amiA* Δ*amiC* Δ*tatABC*) was used for signal peptide library screening and for SDS growth tests with signal peptide-AmiA fusion proteins (27). Strain DADE (as MC4100, Δ*tatABCD* Δ*tatE*; 58) was used for SufI transport assays and DADE-P (as DADE, *pcnB1 zad*-981::Tn*10*d (Kan^r^; 59) was used for co-purification experiments.

All plasmids used and constructed in this study are given in Table S2. Point mutations in plasmids were introduced by Quickchange site-directed mutagenesis (Stratagene) using the primers listed in Table S3. Plasmid pTAT101 was used for low-level production of TatA, TatB and TatC (37). Plasmid pSUSufIss-mAmiA was used to produce SufIss-AmiA, where the SufI signal peptide is fused to the mature portion of AmiA (27). Plasmid pQE80-SufIhis was used to produce his-tagged SufI (27).

Plasmids pSUDsbAss-mAmiA, pSUDsbAssi16K-mAmiA (with an additional lysine codon inserted after codon 16 of the DbsA signal peptide), pSUOmpAss-mAmiA, and pSUOmpAssi18K-mAmiA (with an additional lysine codon inserted after codon 18 of the OmpA signal peptide) were constructed according to (27). Briefly, DNA fragments encoding DsbAss, DsbAssi16K, OmpAss, and OmpAssi18K were amplified by PCR using MC4100 genomic DNA as template, using primer pairs DsbAss-FE / DsbAss-R, DsbAss-FE / DsbAss16inK-R, OmpA-FE / OmpAss-R, and OmpA-FE / OmpA18inK-R, respectively. DNA fragments encoding the corresponding mature domain of AmiA were amplified by PCR using MC4100 genomic DNA as template with primer pairs OmpA-mAmiA-F / amiA-mRX, or DsbA-mAmiA-F / amiA-mRX. The DNA fragments encoding the signal peptides and the mature domain of AmiA were fused by overlap extension PCR, giving DNA fragments DsbAss-mAmiA, DsbAssi16K-mAmiA, OmpAss-mAmiA, and OmpAi18Kss-mAmiA, which were finally cloned into pSU18 vector following digestion with *Eco*RI and *Xba*I.

Plasmid pFAT75BC-SufIFLAG was modified from pFAT75ΔA-SufIhis (18) via quickchange using primers FAT75SufIFLAG-1 / FAT75SufIFLAG-2. Plasmid pFATBChis-SufIFLAG was modified from pFATSufIFLAG via quickchange using primers FAT75TatChis-1 / FAT75TatChis-2. Plasmid pFATBChis-OmpAssi18KSufIFLAG has the SufI signal peptide coding region substituted for DNA encoding OmpAssi18K, and was constructed using a restriction enzyme-free cloning method according to (60). Briefly a DNA fragment covering OmpAssi18K was PCR amplified using pSUOmpAssi18K-mAmiA as template, with primer pair FATHF-OmpA-F / FATHF-OmpA18K-R. The resultant DNA fragment was used as a primer to amplify the whole pFATBChis-SufIFLAG plasmid using the PCR program: 95°C 2min followed by 15 cycles of 95 °C 30s, 48 °C 1.5 min and 68 °C 15 min and a final extension at 68 °C for 10 min. The PCR product was subject to *Dpn*I digestion and introduced into *E. coli* JM109 competent cells by transformation. The resultant plasmid was verified by DNA sequencing. Plasmids pQEBChis-OmpAFLAG and pQEBChis-DsbAFLAG were used for co-production of TatB, his-tagged TatC and FLAG-tagged OmpA or DsbA, respectively, and were constructed as follows. A DNA fragment encoding TatB and his-tagged TatC was amplified using pFATBChis-SufIFLAG as template with primer pair QEF / FAT75TatChis-2, and was ligated, via an *Apa*I restriction site, to a DNA fragment encoding FLAG-tagged OmpA (that was amplified using MC4100 genomic DNA as template with primer pair FATHF-OmpA-F / OmpAFLAG-SR) or FLAG-tagged DsbA (amplified similarly using primer pair FATHF-OmpA-F / DsbAFLAG-SR). The ligated fragment was enriched by using the ligation mixture as a template in a PCR reaction with primer pair QEF / OmpAFLAG-SR, or QEF / DsbAAFLAG-SR, presectively. The amplified fragment was gel-purified, digested with *Eco*RI and *Sal*I and cloned into similarly-digested pQE80. Constructs were verified by DNA sequencing.

### Mutant library construction and screening

To construct a random library of substitutions at codon 8 of the SufI signal peptide, site-directed mutagenesis was carried out via Quickchange using a pair of random primers SufIF8X1 and SufIF8X2 (Table S3) and pSUSufIss-mAmiA as template. The PCR product was subsequently introduced into XL1-Gold ultra-competent cells (Agilent). Transformants were scraped from plates, resuspended in LB, pooled and cultured overnight after which plasmid DNA was isolated and taken as the F8X random library. The library contained approximately 5 000 clones and random sequencing of eight of them revealed substitutions of the TTC codon to CTT, ATC GGT, GTG, GGT, CAA, AGA and GGG.

The signal peptide mutagenesis library in plasmid pSUSufIss-mAmiA was constructed as described (27). Briefly, an error-containing DNA fragment covering the *sufI* signal sequence was amplified by error-prone PCR using primers SufIF and SufIR and pSUSufIss-mAmiA as template. This fragment was used as a megaprimer to amplify the whole pSUSufIss-mAmiA plasmid. The amplified plasmid was introduced into XL1-gold ultra-competent cells following nick repair using T4 polynucleotide kinase and T4 ligase. Transformants were scraped from plates, resuspended in LB, pooled and used to inoculate fresh LB to an initial OD_600_ of 0.2. Cells were grown at 37 °C until OD_600_ reached 2 after which plasmid DNA was prepared and taken as the signal peptide mutagenesis library.

For library screening, plasmid pTAT101 harboring the *tatC* point substitution of interest (along with wild type *tatAB*) was introduced into MC4100 Δ*amiA* Δ*amiC* Δ*tatABC*. Subsequently the mutant library was introduced and cells were plated onto LB agar containing 2% SDS. Plasmids were isolated from colonies growing on this selective medium and mutations identified by sequencing.

### Protein methods

Co-purification of TatBC-substrate complexes was carried out as described (27). Briefly, an overnight culture of DADE-P harboring plasmid pFATBChis-SufIFLAG or its derivatives was subcultured in LB supplemented with 0.5% glycerol and appropriate antibiotics for 2.5 hours at 37 °C with shaking. Following supplemented with 0.4 mM isopropyl-β-D-galactopyranoside (IPTG), cells were incubated overnight at 30 °C. The following morning, cells were harvested, resuspended in 200μL of 2X lysis buffer (100 mM NaH_2_PO_4_ pH 8.0, 600 mM NaCl, 40 mM imidazole, 50 mg lysozyme, DNase I, and protease inhibitor), and mixed gently at room temperature for one hour. Cells were then frozen at -80 °C for 1 hour and thawed at room temperature. An equal volume of 2.5% digitonin was added to the cells and the samples were solubilized for 1 hour at 4°C. The insoluble material was pelleted by centrifugation at 4 °C. A 30 μL of the supernatant was mixed with 2x Laemmli buffer which was taken as the input sample and the remaining supernatant was mixed with 50 μL wash buffer-equilibrated nickel beads (Profinity™ IMAC Ni-Charged Resin, Bio-Rad, catalog number 156-0131) for one hour. The nickel beads were pelleted, washed three times with 1 mL wash buffer (50 mM NaH_2_PO_4_, pH 8.0, 300 mM NaCl, 40 mM imidazole, 0.03 % digitonin) and then mixed with 100 μL elution buffer (50 mM NaH_2_PO_4_, pH 8.0, 300 mM NaCl, 700 mM imidazole, 0.03 % digitonin). The beads were incubated for 10 min with shaking and then pelleted. The supernatant (elution fraction) was taken, mixed with an equal volume of 2x Laemmli buffer, and 20 μl of the sample was subjected to SDS-PAGE followed by western-blotting with anti-His (Anti-6X His tag® antibody [GT359] (HRP), Abcam, catalog number ab184607), anti-TatB (61), or anti-FLAG antibodies (Monoclonal ANTI-FLAG® M2 antibody produced in mouse. Sigma catalog number F1804). Secondary antibody was goat anti-rabbit IgG (HRP Conjugate, Bio-Rad, catalog number 170-6515) or goat anti-mouse IgG (HRP Conjugate, Bio-Rad, catalog number 1706516).

Subcellular fractionation was carried out as described previously (27). Briefly, overnight cultures of strain DADE harboring pTAT101 or the cognate empty vector pTH19kr along with pQE80-SufIhis or its derivatives were subcultured at 1:50 in LB supplemented with 1mM IPTG and grown at 37 °C until OD_600_ reached 1. Where the RR-KK substitution was present in the SufI, no IPTG was used (as for unknown reasons this substitution results in high level expression of SufI even in the absence of IPTG). For whole cell samples, cells were pelleted from 5 mL of the culture, resuspended in 250 μL resuspension buffer (50mM Tris-HCl, pH 7.6, 2 mM EDTA), and lysed by sonicating for 15 s. The cell lysate was mixed with an equal volume of 2x Laemmli buffer and boiled at 95 °C for 10 min. For preparation of periplasm, cells were pelleted from 20 mL of the culture and resuspended in 500 μL fractionation buffer (20mM Tris-HCl, pH 7.6, 2 mM EDTA, 20% sucrose (w/v)). 0.6 mg/mL freshly made lysozyme was added and the cells were incubated at room temperature for 20 min. The cells were then pelleted by centrifugation and the supernatant was taken and mixed with equal volume of 2x Laemmli buffer. Aliquots (20 μL) of the whole cell or periplasmic fraction samples were separated by SDS-PAGE (10% acrylamide) followed by western blotting with anti-His, or anti-his and anti-RNA polymerase β-subunit mixed antibodies.

### Prediction of Sec signal peptides

All of the protein sequences encoded by *E. coli* MG1655 were analyzed using the SignalP 4.1 server (http://www.cbs.dtu.dk/services/SignalP; (55)) with parameters ‘Gram-negative bacteria’ and ‘input sequence do not include TM regions’ selected. Inner membrane proteins were removed manually from the output Sec substrate candidates.

## Acknowledgements

This work was supported by the China Scholarship Council (through a studentship to QH). TP is a Wellcome Trust Investigator.

---

**Table S1.**
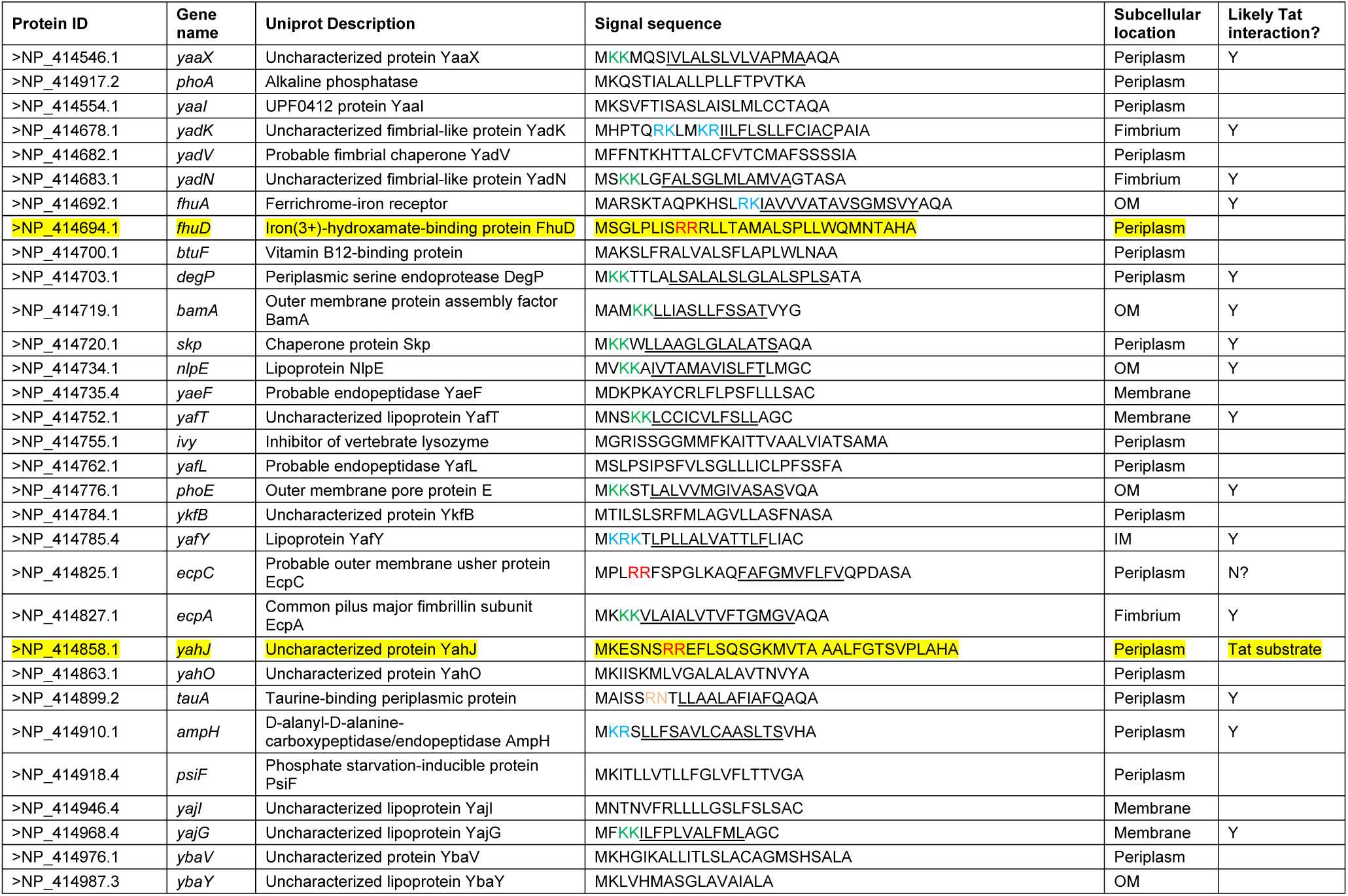
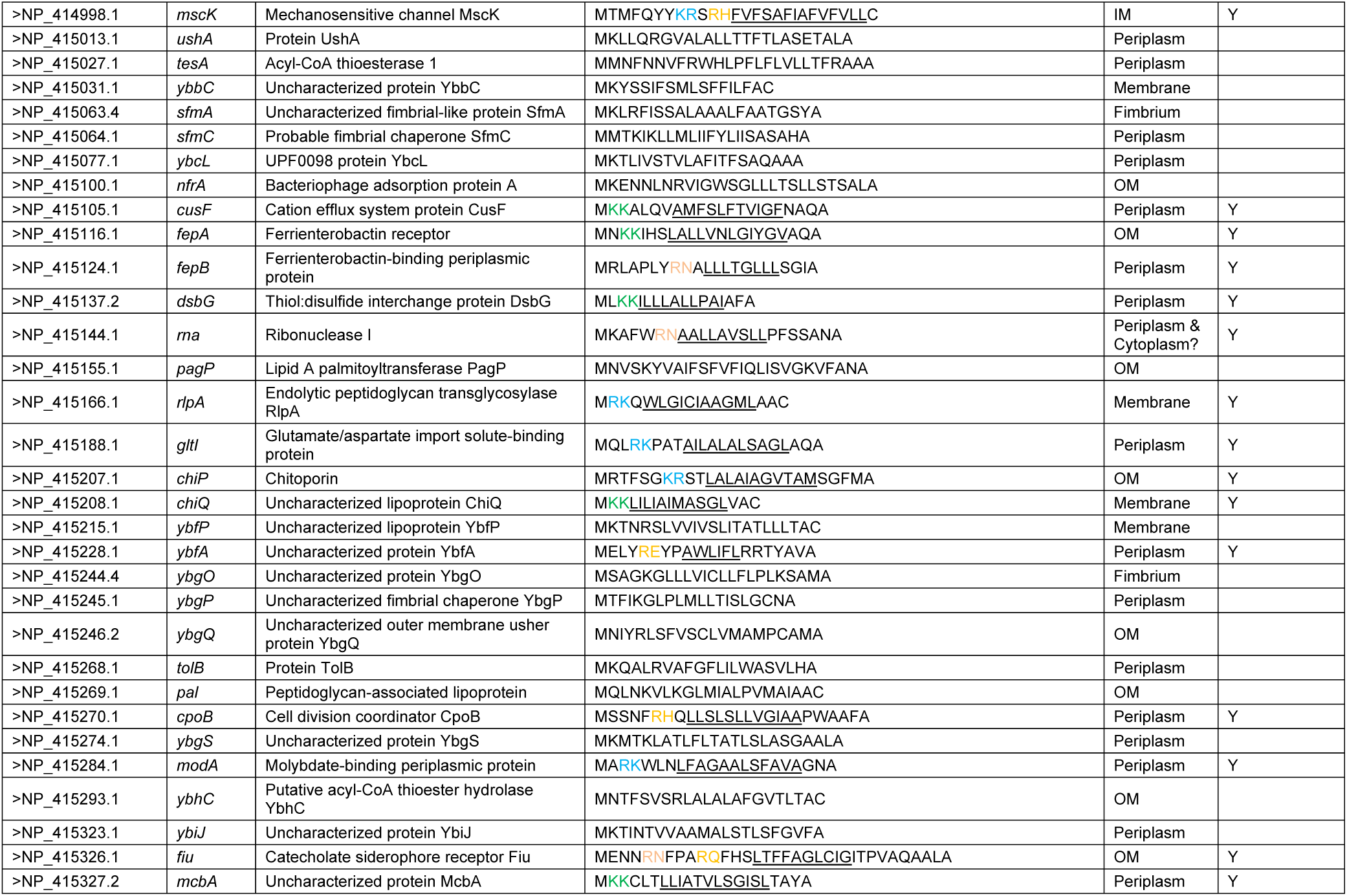
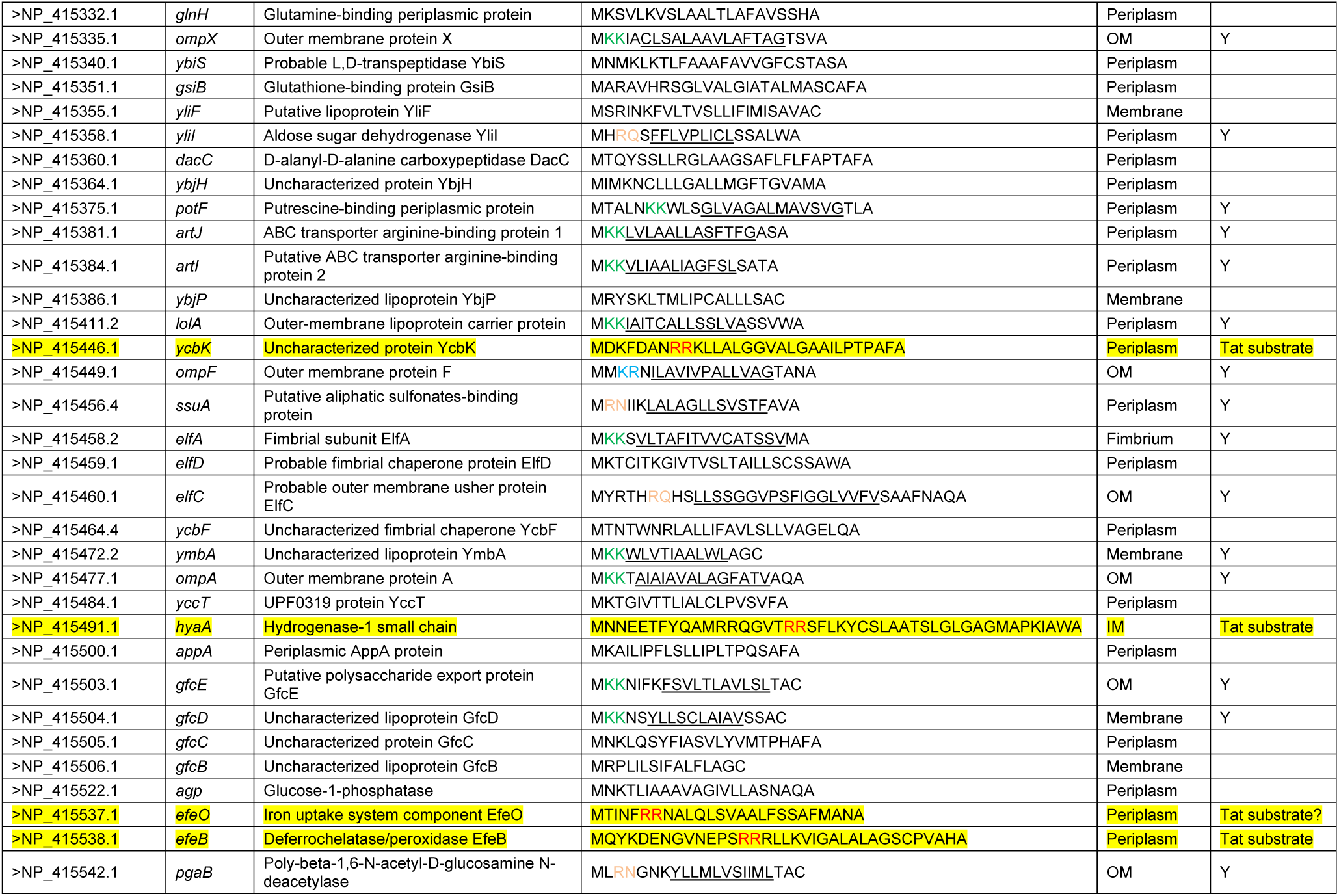
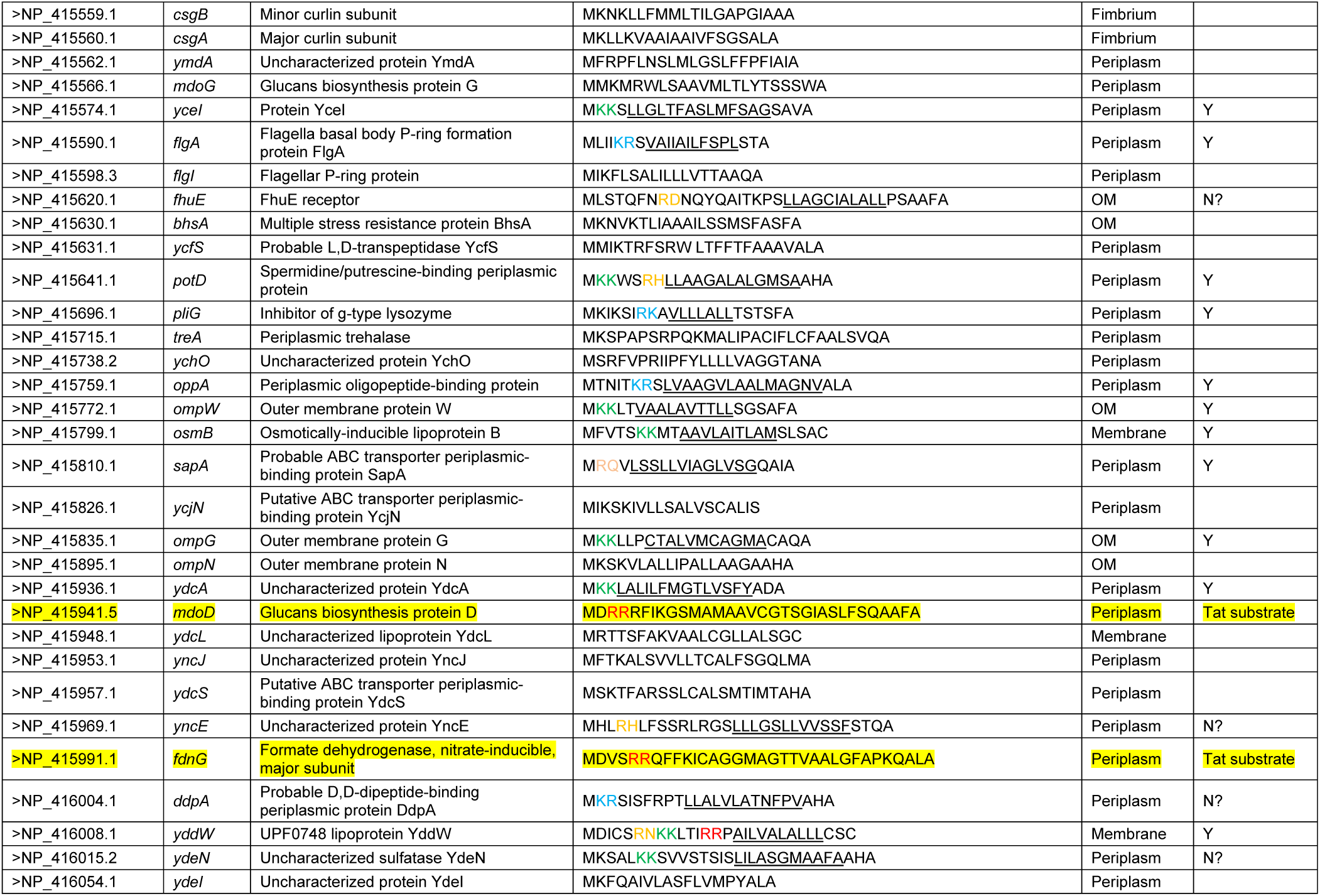
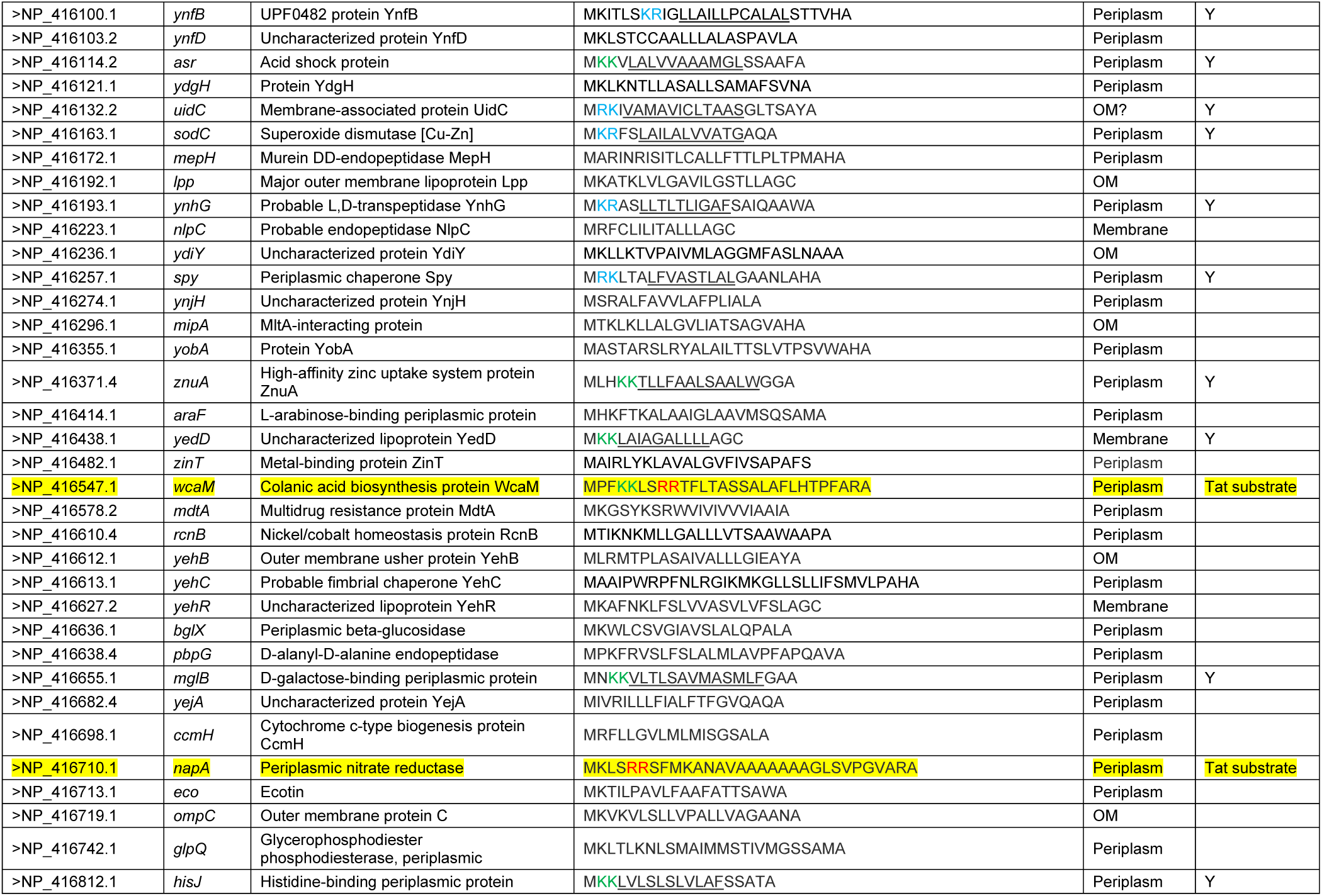
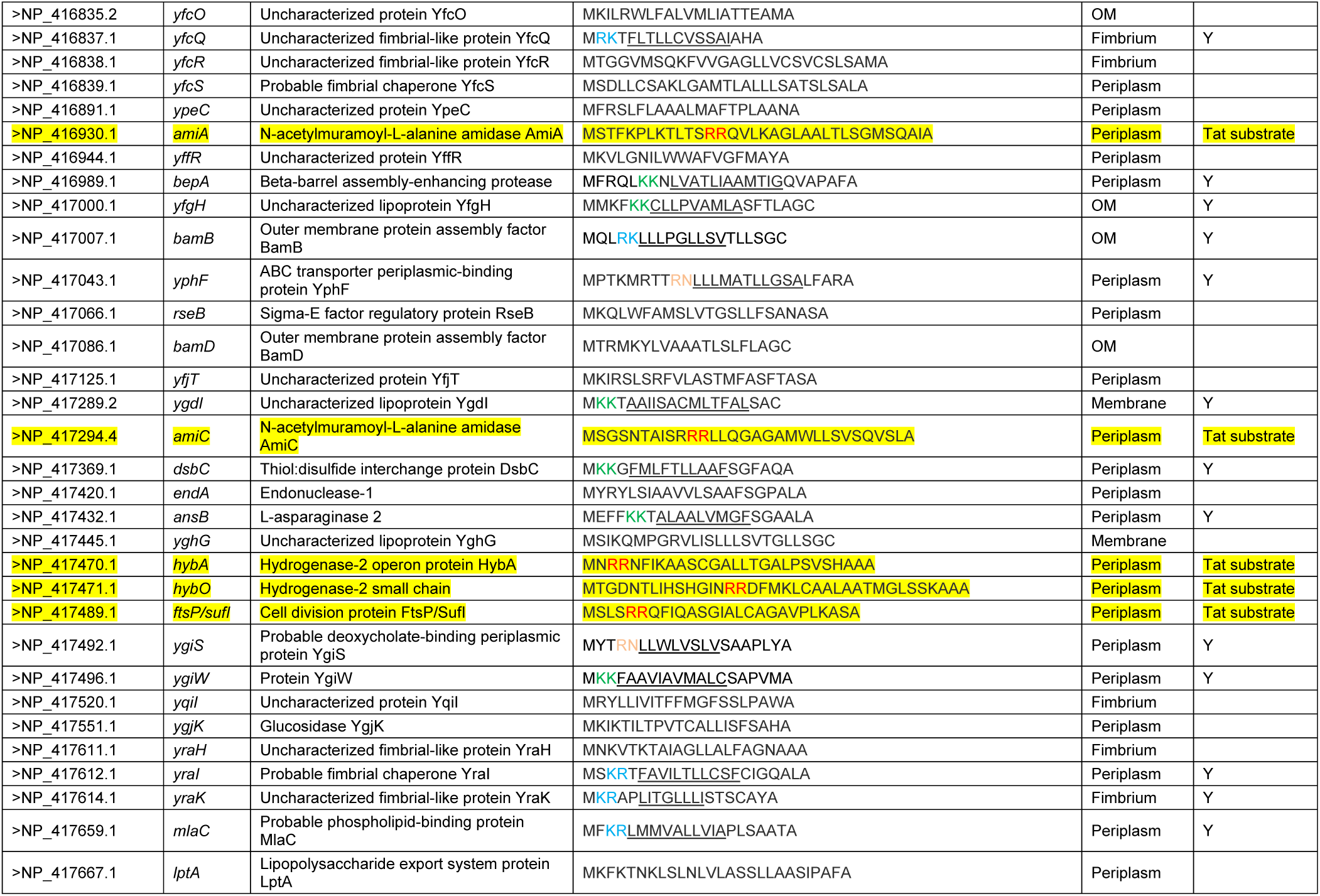
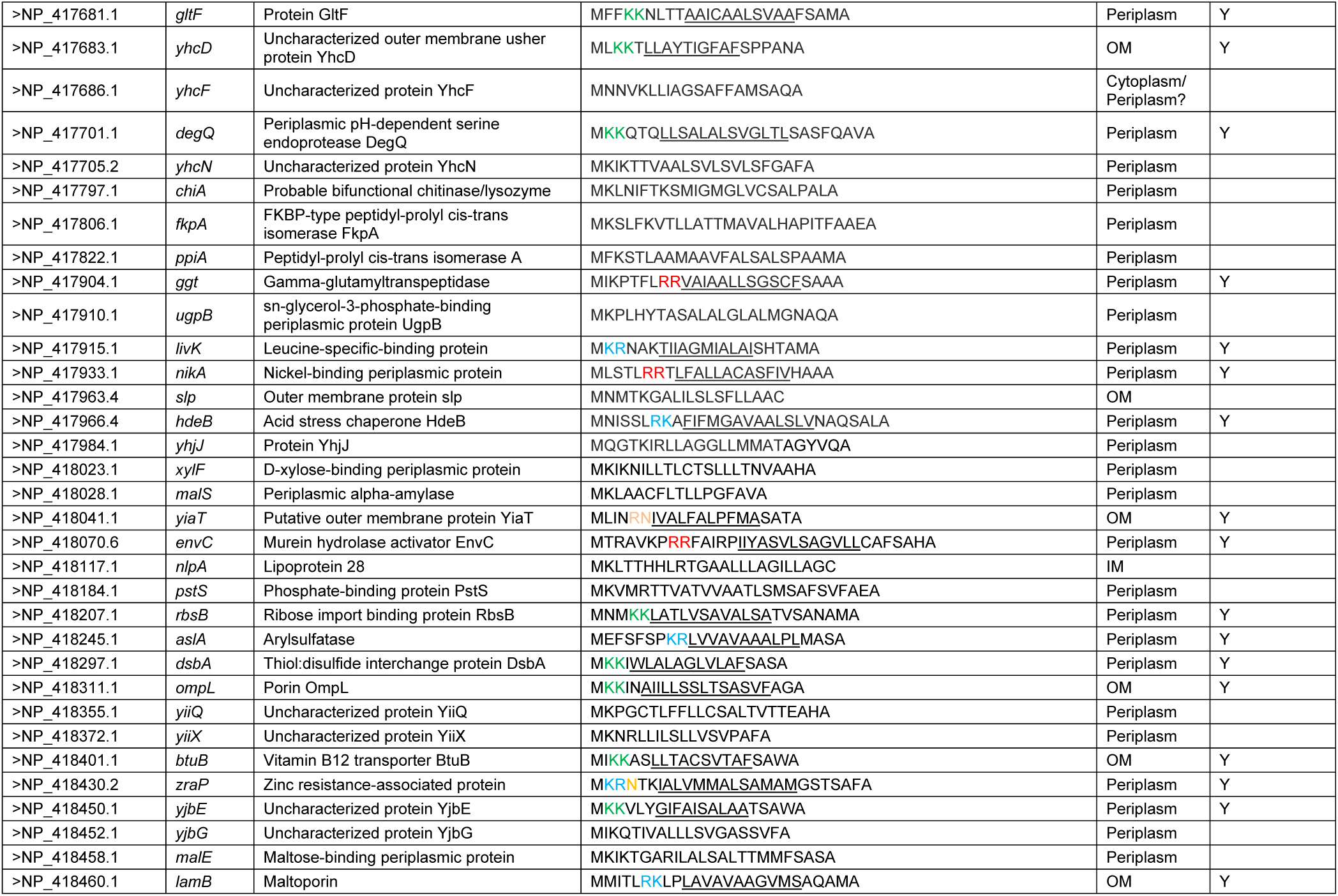
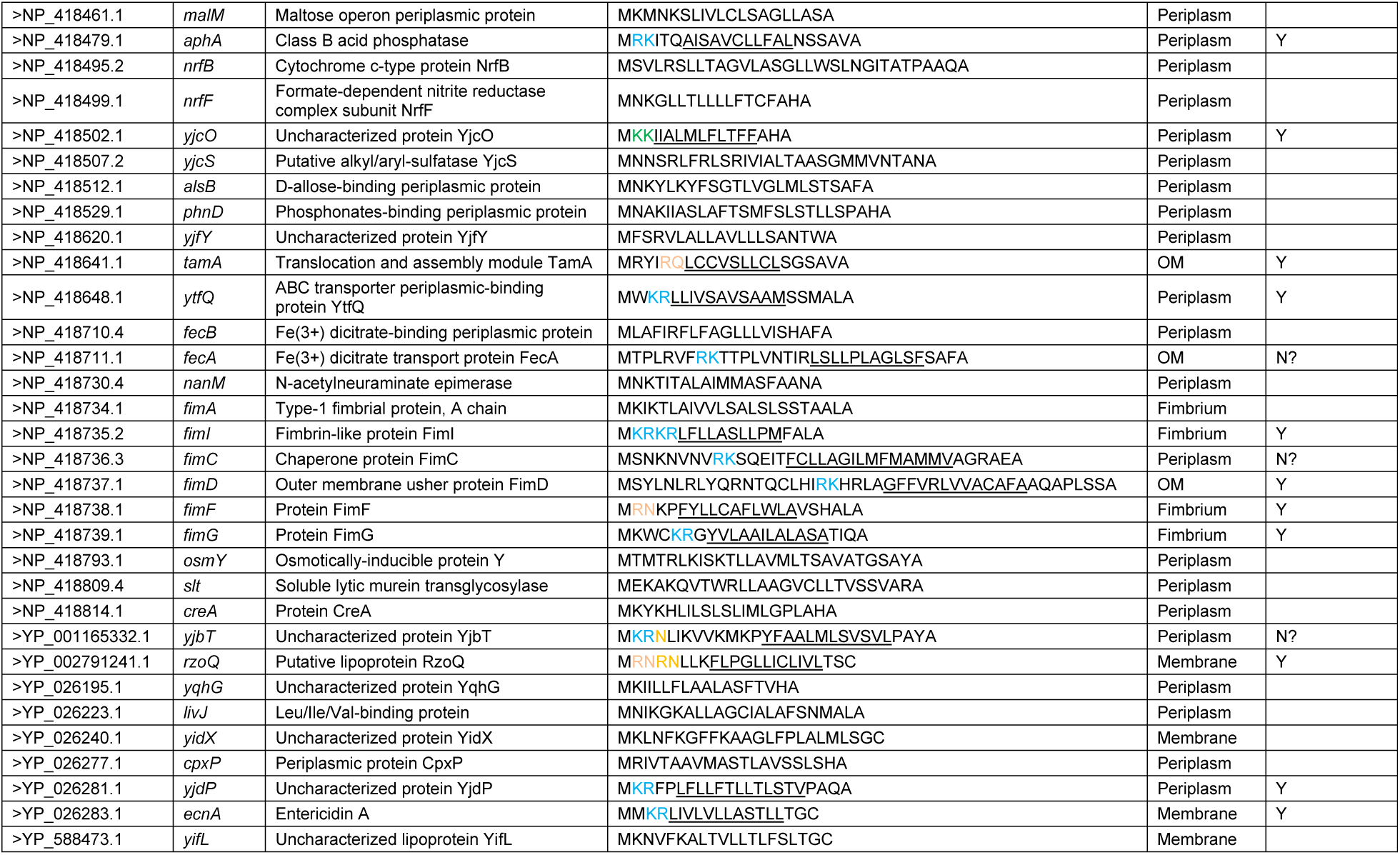
*E. coli* proteins with known or predicted signal peptides. Protein sequences encoded by *E. coli* MG1655 were retrieved from Uniprot (www.uniprot.org) and analyzed for the presence of a signal peptide using the SignalP 4.1 server (http://www.cbs.dtu.dk/services/SignalP). The n-regions were analyzed manually; twin arginines are colored red, twin lysines green, lys-arg or arg-lys are shown in blue. Pairings of Arg-Asn, Arg-Asp, Arg-Gln, Arg-Glu and Arg-His (which from this study can be suppressed by increased signal peptide hydrophobicity) are shown in orange. For those signal peptides harboring any of these dipeptides in their n-regions the likely location of the h-region is shown in underline. Pairings within five residues of the h-region were considered likely to interact with the Tat pathway. Known or probable Tat substrates are shown in yellow highlight.

**Table S2.**
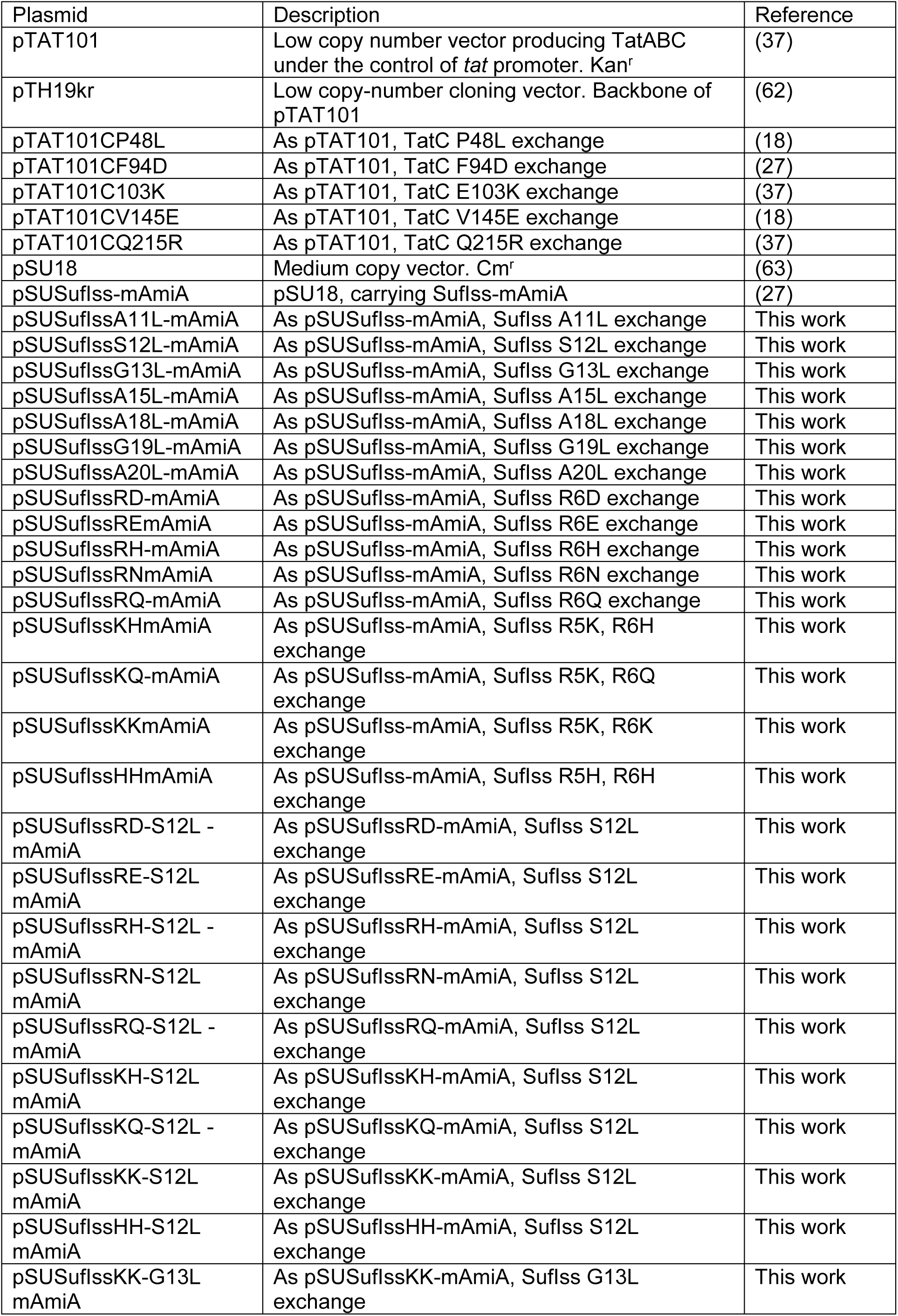
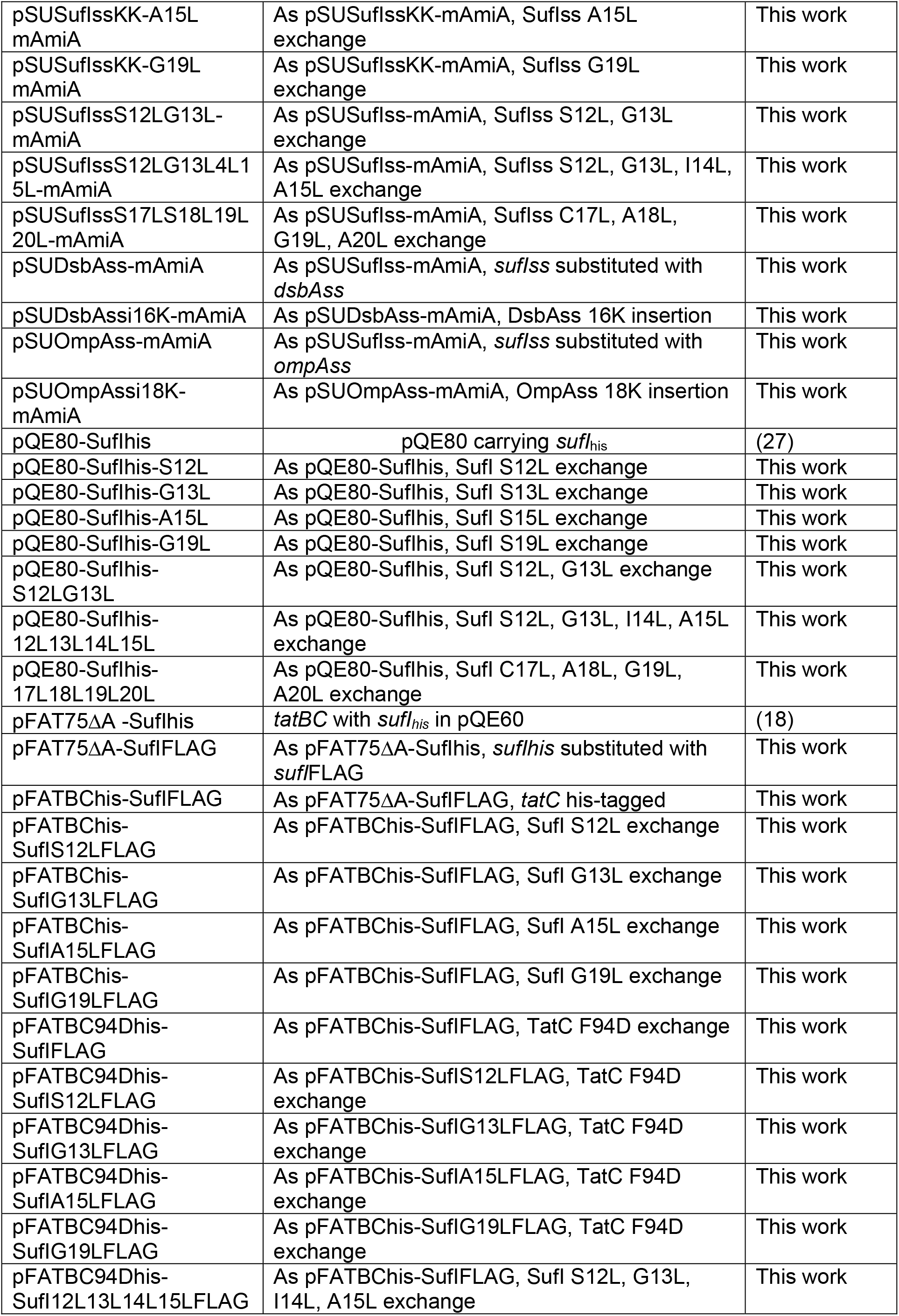
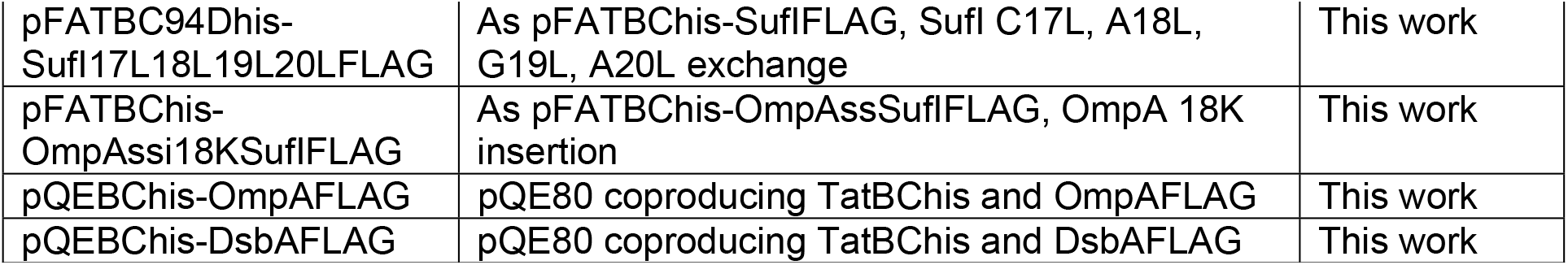
Plasmids used and constructed in this study.

| Plasmid | Description | Reference |
| --- | --- | --- |
| pTAT101 | Low copy number vector producing TatABC under the control of *tat* promoter. Kan<sup>r</sup> | (37) |
| pTH19kr | Low copy-number cloning vector. Backbone of pTAT101 | (62) |
| pTAT101CP48L | As pTAT101, TatC P48L exchange | (18) |
| pTAT101CF94D | As pTAT101, TatC F94D exchange | (27) |
| pTAT101C103K | As pTAT101, TatC E103K exchange | (37) |
| pTAT101CV145E | As pTAT101, TatC V145E exchange | (18) |
| pTAT101CQ215R | As pTAT101, TatC Q215R exchange | (37) |
| pSU18 | Medium copy vector. Cm<sup>r</sup> | (63) |
| pSUSufI<sup>ss</sup>-mAmiA | pSU18, carrying SufI<sup>ss</sup>-mAmiA | (27) |
| pSUSufI<sup>ss</sup>A11L-mAmiA | As pSUSufI<sup>ss</sup>-mAmiA, SufI<sup>ss</sup> A11L exchange | This work |
| pSUSufI<sup>ss</sup>S12L-mAmiA | As pSUSufI<sup>ss</sup>-mAmiA, SufI<sup>ss</sup> S12L exchange | This work |
| pSUSufI<sup>ss</sup>G13L-mAmiA | As pSUSufI<sup>ss</sup>-mAmiA, SufI<sup>ss</sup> G13L exchange | This work |
| pSUSufI<sup>ss</sup>A15L-mAmiA | As pSUSufI<sup>ss</sup>-mAmiA, SufI<sup>ss</sup> A15L exchange | This work |
| pSUSufI<sup>ss</sup>A18L-mAmiA | As pSUSufI<sup>ss</sup>-mAmiA, SufI<sup>ss</sup> A18L exchange | This work |
| pSUSufI<sup>ss</sup>G19L-mAmiA | As pSUSufI<sup>ss</sup>-mAmiA, SufI<sup>ss</sup> G19L exchange | This work |
| pSUSufI<sup>ss</sup>A20L-mAmiA | As pSUSufI<sup>ss</sup>-mAmiA, SufI<sup>ss</sup> A20L exchange | This work |
| pSUSufI<sup>ss</sup>RD-mAmiA | As pSUSufI<sup>ss</sup>-mAmiA, SufI<sup>ss</sup> R6D exchange | This work |
| pSUSufI<sup>ss</sup>REmAmiA | As pSUSufI<sup>ss</sup>-mAmiA, SufI<sup>ss</sup> R6E exchange | This work |
| pSUSufI<sup>ss</sup>RH-mAmiA | As pSUSufI<sup>ss</sup>-mAmiA, SufI<sup>ss</sup> R6H exchange | This work |
| pSUSufI<sup>ss</sup>RNmAmiA | As pSUSufI<sup>ss</sup>-mAmiA, SufI<sup>ss</sup> R6N exchange | This work |
| pSUSufI<sup>ss</sup>RQ-mAmiA | As pSUSufI<sup>ss</sup>-mAmiA, SufI<sup>ss</sup> R6Q exchange | This work |
| pSUSufI<sup>ss</sup>KHmAmiA | As pSUSufI<sup>ss</sup>-mAmiA, SufI<sup>ss</sup> R5K, R6H exchange | This work |
| pSUSufI<sup>ss</sup>KQ-mAmiA | As pSUSufI<sup>ss</sup>-mAmiA, SufI<sup>ss</sup> R5K, R6Q exchange | This work |
| pSUSufI<sup>ss</sup>KKmAmiA | As pSUSufI<sup>ss</sup>-mAmiA, SufI<sup>ss</sup> R5K, R6K exchange | This work |
| pSUSufI<sup>ss</sup>HHmAmiA | As pSUSufI<sup>ss</sup>-mAmiA, SufI<sup>ss</sup> R5H, R6H exchange | This work |
| pSUSufI<sup>ss</sup>RD-S12L - mAmiA | As pSUSufI<sup>ss</sup>RD-mAmiA, SufI<sup>ss</sup> S12L exchange | This work |
| pSUSufI<sup>ss</sup>RE-S12L mAmiA | As pSUSufI<sup>ss</sup>RE-mAmiA, SufI<sup>ss</sup> S12L exchange | This work |
| pSUSufI<sup>ss</sup>RH-S12L - mAmiA | As pSUSufI<sup>ss</sup>RH-mAmiA, SufI<sup>ss</sup> S12L exchange | This work |
| pSUSufI<sup>ss</sup>RN-S12L mAmiA | As pSUSufI<sup>ss</sup>RN-mAmiA, SufI<sup>ss</sup> S12L exchange | This work |
| pSUSufI<sup>ss</sup>RQ-S12L - mAmiA | As pSUSufI<sup>ss</sup>RQ-mAmiA, SufI<sup>ss</sup> S12L exchange | This work |
| pSUSufI<sup>ss</sup>KH-S12L mAmiA | As pSUSufI<sup>ss</sup>KH-mAmiA, SufI<sup>ss</sup> S12L exchange | This work |
| pSUSufI<sup>ss</sup>KQ-S12L - mAmiA | As pSUSufI<sup>ss</sup>KQ-mAmiA, SufI<sup>ss</sup> S12L exchange | This work |
| pSUSufI<sup>ss</sup>KK-S12L mAmiA | As pSUSufI<sup>ss</sup>KK-mAmiA, SufI<sup>ss</sup> S12L exchange | This work |
| pSUSufI<sup>ss</sup>HH-S12L mAmiA | As pSUSufI<sup>ss</sup>HH-mAmiA, SufI<sup>ss</sup> S12L exchange | This work |
| pSUSufI<sup>ss</sup>KK-G13L mAmiA | As pSUSufI<sup>ss</sup>KK-mAmiA, SufI<sup>ss</sup> G13L exchange | This work |
| pSUSufI<sub>ss</sub>KK-A15L-mAmiA | As pSUSufI<sub>ss</sub>KK-mAmiA, SufI<sub>ss</sub> A15L exchange | This work |
| pSUSufI<sub>ss</sub>KK-G19L-mAmiA | As pSUSufI<sub>ss</sub>KK-mAmiA, SufI<sub>ss</sub> G19L exchange | This work |
| pSUSufI<sub>ss</sub>S12LG13L-mAmiA | As pSUSufI<sub>ss</sub>-mAmiA, SufI<sub>ss</sub> S12L, G13L exchange | This work |
| pSUSufI<sub>ss</sub>S12LG13L4L15L-mAmiA | As pSUSufI<sub>ss</sub>-mAmiA, SufI<sub>ss</sub> S12L, G13L, I14L, A15L exchange | This work |
| pSUSufI<sub>ss</sub>S17LS18L19L20L-mAmiA | As pSUSufI<sub>ss</sub>-mAmiA, SufI<sub>ss</sub> C17L, A18L, G19L, A20L exchange | This work |
| pSUDsbAss-mAmiA | As pSUSufI<sub>ss</sub>-mAmiA, *sufI<sub>ss</sub>* substituted with *dsbAss* | This work |
| pSUDsbAssi16K-mAmiA | As pSUDsbAss-mAmiA, DsbAss 16K insertion | This work |
| pSUOmpAss-mAmiA | As pSUSufI<sub>ss</sub>-mAmiA, *sufI<sub>ss</sub>* substituted with *ompAss* | This work |
| pSUOmpAssi18K-mAmiA | As pSUOmpAss-mAmiA, OmpAss 18K insertion | This work |
| pQE80-SufI<sub>his</sub> | pQE80 carrying *sufI<sub>his</sub>* | (27) |
| pQE80-SufI<sub>his</sub>-S12L | As pQE80-SufI<sub>his</sub>, SufI S12L exchange | This work |
| pQE80-SufI<sub>his</sub>-G13L | As pQE80-SufI<sub>his</sub>, SufI S13L exchange | This work |
| pQE80-SufI<sub>his</sub>-A15L | As pQE80-SufI<sub>his</sub>, SufI S15L exchange | This work |
| pQE80-SufI<sub>his</sub>-G19L | As pQE80-SufI<sub>his</sub>, SufI S19L exchange | This work |
| pQE80-SufI<sub>his</sub>-S12LG13L | As pQE80-SufI<sub>his</sub>, SufI S12L, G13L exchange | This work |
| pQE80-SufI<sub>his</sub>-12L13L14L15L | As pQE80-SufI<sub>his</sub>, SufI S12L, G13L, I14L, A15L exchange | This work |
| pQE80-SufI<sub>his</sub>-17L18L19L20L | As pQE80-SufI<sub>his</sub>, SufI C17L, A18L, G19L, A20L exchange | This work |
| pFAT75ΔA-SufI<sub>his</sub> | *tatBC* with *sufI<sub>his</sub>* in pQE60 | (18) |
| pFAT75ΔA-SufIFLAG | As pFAT75ΔA-SufI<sub>his</sub>, *sufI<sub>his</sub>* substituted with *sufIFLAG* | This work |
| pFATBChis-SufIFLAG | As pFAT75ΔA-SufIFLAG, *tatC* his-tagged | This work |
| pFATBChis-SufIS12LFLAG | As pFATBChis-SufIFLAG, SufI S12L exchange | This work |
| pFATBChis-SufIG13LFLAG | As pFATBChis-SufIFLAG, SufI G13L exchange | This work |
| pFATBChis-SufIA15LFLAG | As pFATBChis-SufIFLAG, SufI A15L exchange | This work |
| pFATBChis-SufIG19LFLAG | As pFATBChis-SufIFLAG, SufI G19L exchange | This work |
| pFATBC94Dhis-SufIFLAG | As pFATBChis-SufIFLAG, TatC F94D exchange | This work |
| pFATBC94Dhis-SufIS12LFLAG | As pFATBChis-SufIS12LFLAG, TatC F94D exchange | This work |
| pFATBC94Dhis-SufIG13LFLAG | As pFATBChis-SufIG13LFLAG, TatC F94D exchange | This work |
| pFATBC94Dhis-SufIA15LFLAG | As pFATBChis-SufIA15LFLAG, TatC F94D exchange | This work |
| pFATBC94Dhis-SufIG19LFLAG | As pFATBChis-SufIG19LFLAG, TatC F94D exchange | This work |
| pFATBC94Dhis-SufI12L13L14L15LFLAG | As pFATBChis-SufIFLAG, SufI S12L, G13L, I14L, A15L exchange | This work |
| pFATBC94Dhis-Sufl17L18L19L20LFLAG | As pFATBChis-SuflFLAG, Sufl C17L, A18L, G19L, A20L exchange | This work |
| pFATBChis-OmpAssi18KSuflFLAG | As pFATBChis-OmpAssSuflFLAG, OmpA 18K insertion | This work |
| pQEBChis-OmpAFLAG | pQE80 coproducing TatBChis and OmpAFLAG | This work |
| pQEBChis-DsbAFLAG | pQE80 coproducing TatBChis and DsbAFLAG | This work |

**Table S3.**
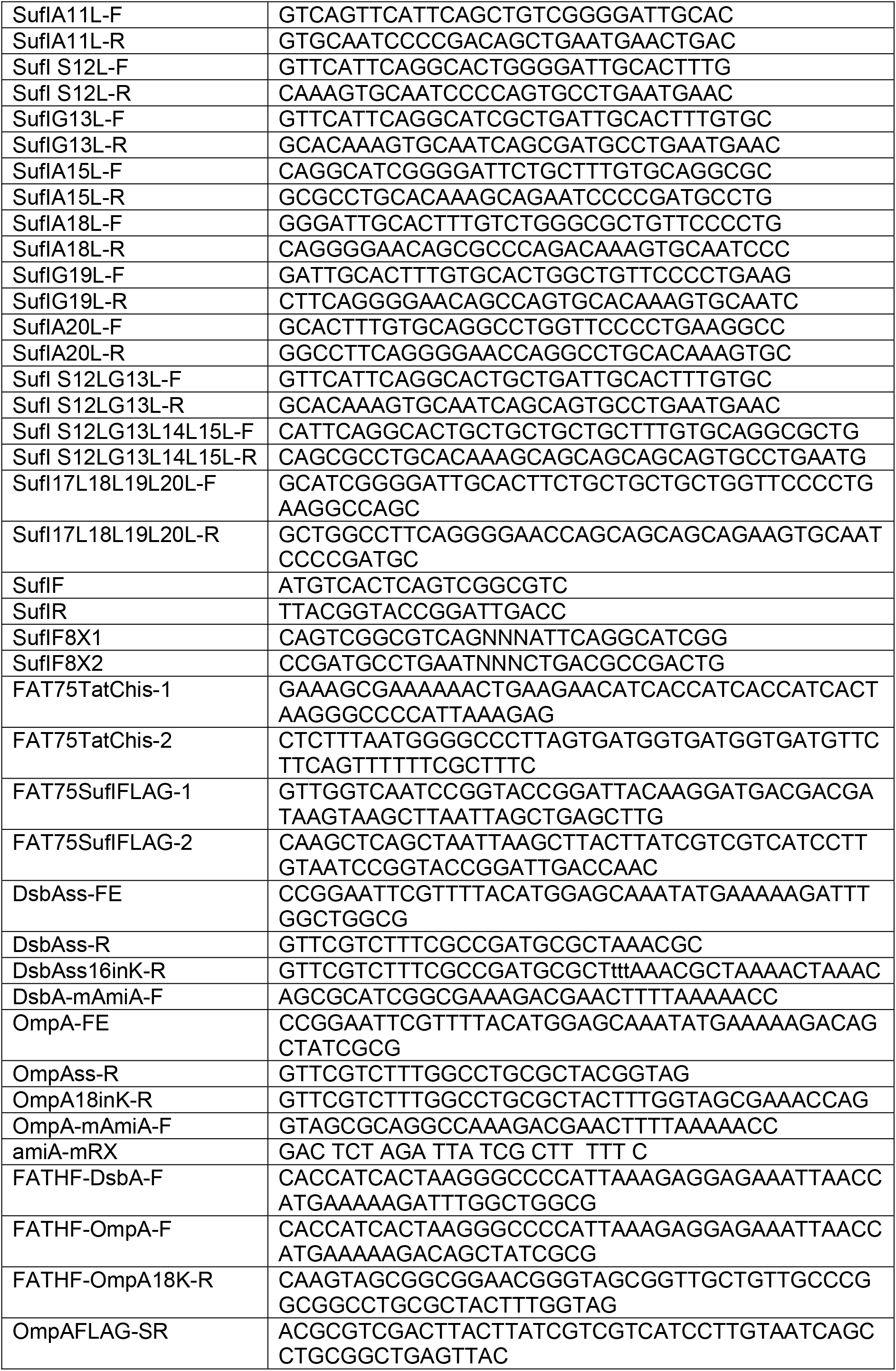
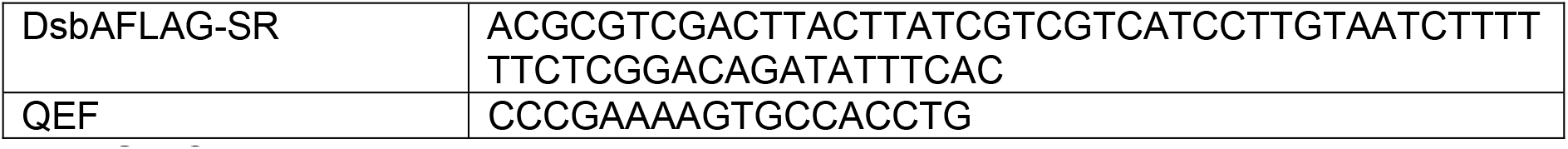
Oligonucleotides used in this study.

**Figure S1.**
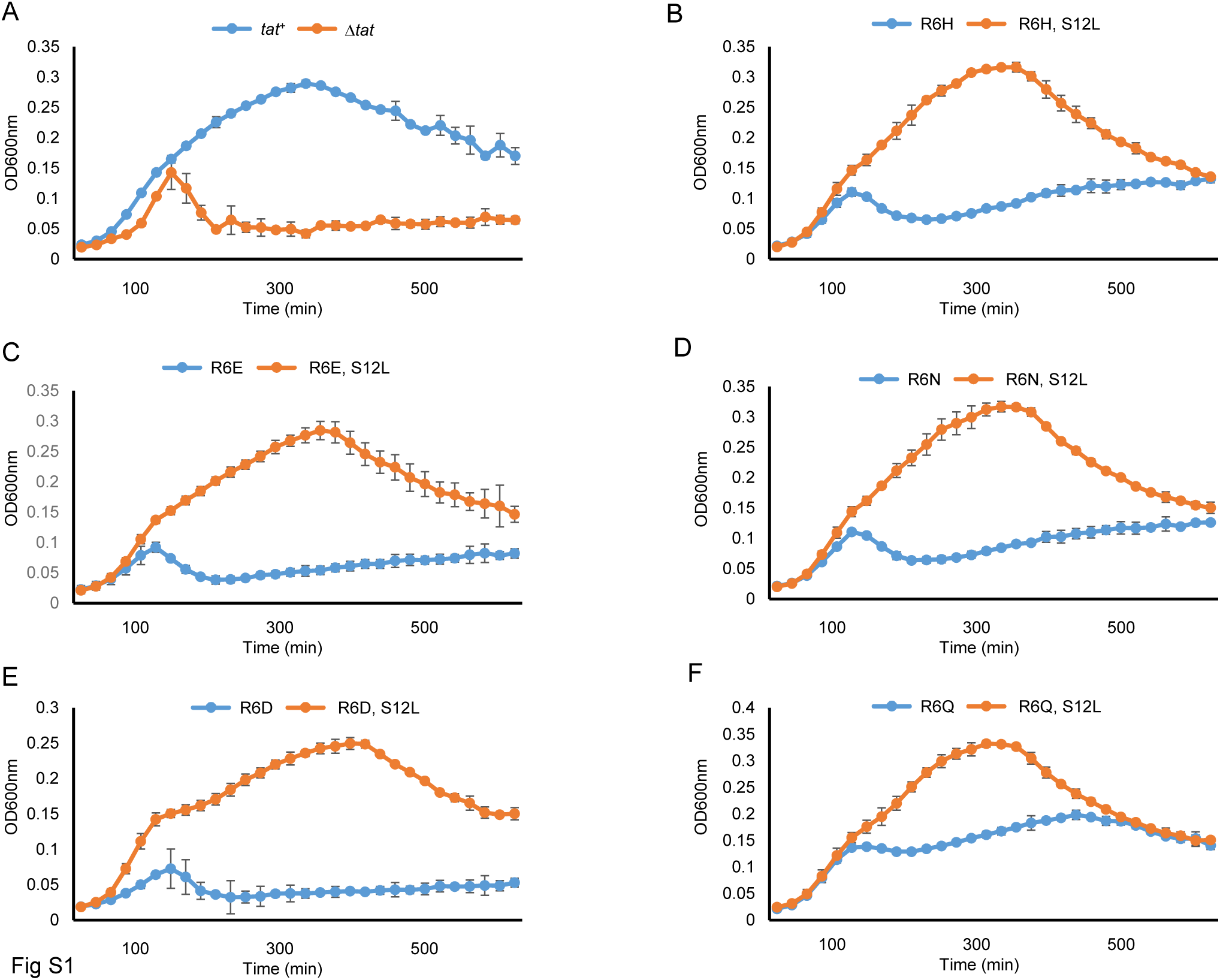
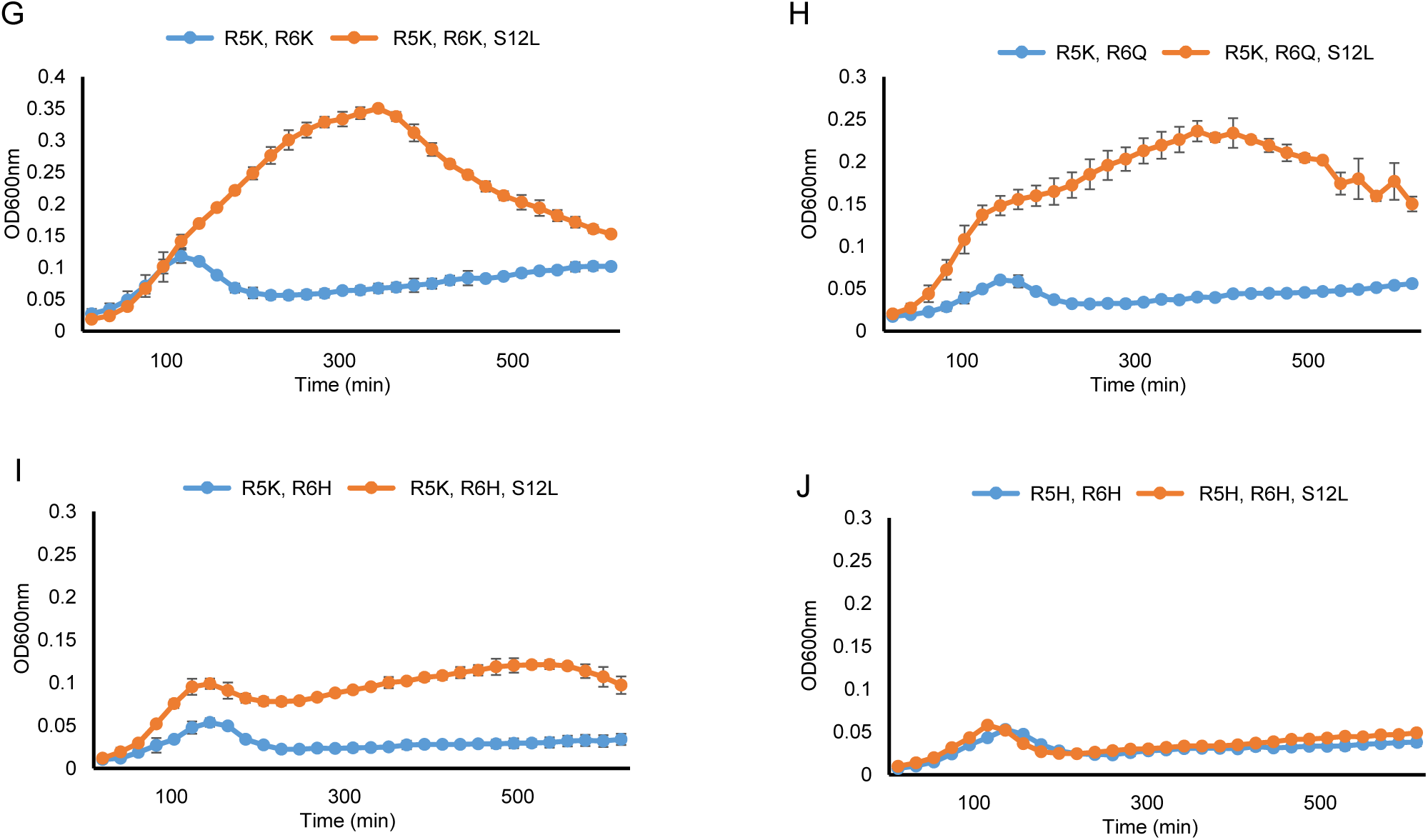
The SufIS12L substitution can act in *cis* to suppress inactivating substitutions in the SufI signal peptide twin-arginine motif. Overnight cultures of strain MC4100 Δ*amiA* Δ*amiC* Δ*tatABC* harboring A. pTH19kr alongside pSUSufIss-mAmiA (Δ*tat*) or pTAT101 (producing wild type TatABC) alongside unsubstituted pSUSufIss-mAmiA (*tat*^+^), or B. – J. pTAT101 (producing wild type TatABC) alongside pSUSufIss-mAmiA encoding the indicated substitutions in the SufI signal peptide were sub-cultured at 1:100 dilution into LB supplemented with 0.5% SDS (final concentration) and grown at 37 °C without shaking The optical density at 600nm was monitored every 20min using a plate reader. Error bars are ± SD, *n*=3 biological replicates.

**Figure S2.**
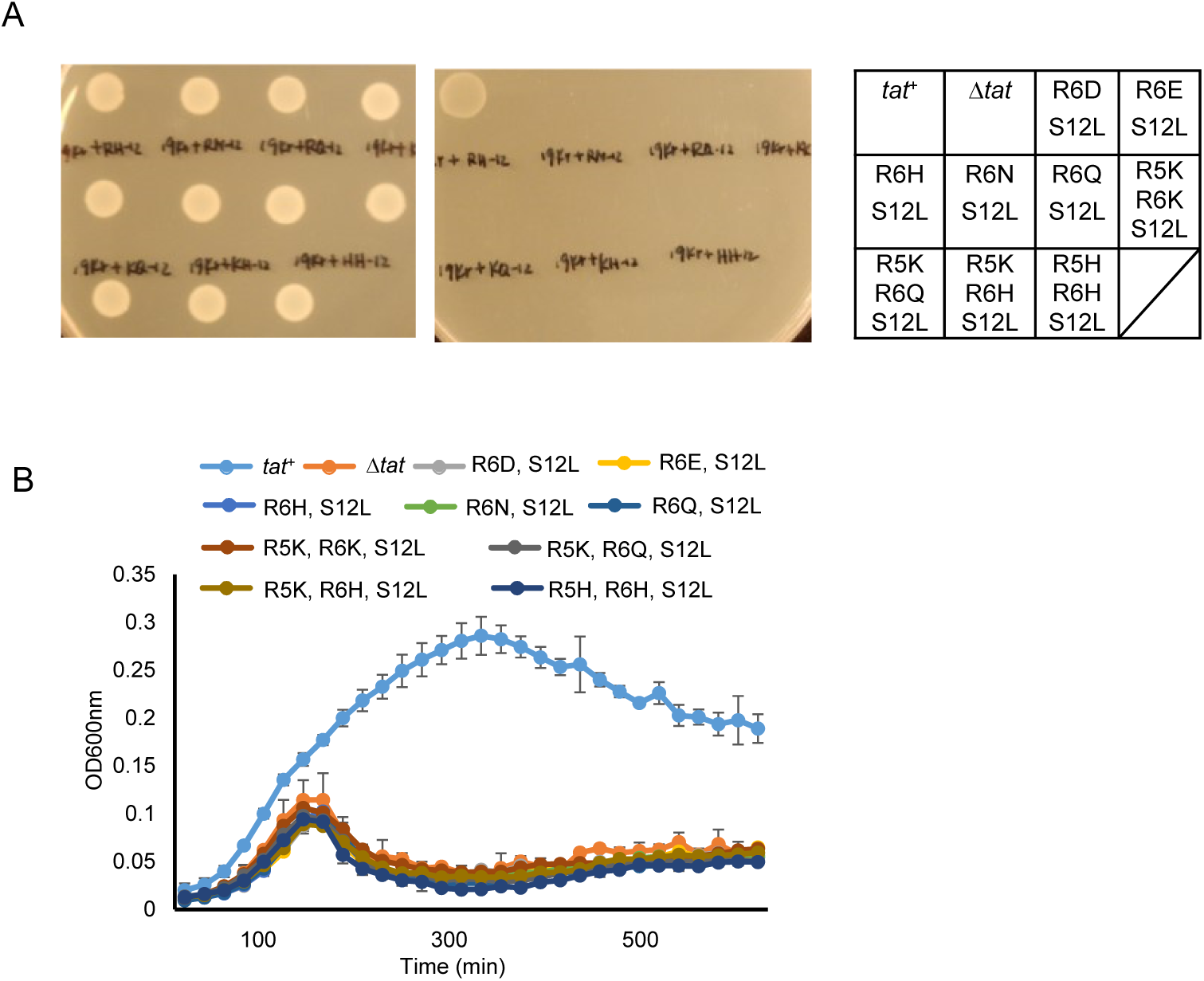
The twin-arginine substituted SufI^S12L^ signal peptide variants mediate strict Tat-dependent transport of AmiA. A and B. Overnight cultures of strain MC4100 Δ*amiA* Δ*amiC* Δ*tatABC* harboring either pTAT101 (producing wild type TatABC) alongside unsubstituted pSUSufIss-mAmiA (*tat*^+^) or pTH19kr alongside either unsubstituted pSUSufIss-mAmiA (Δ*tat*) or pSUSufIss-mAmiA encoding the indicated substitutions in the SufI signal peptide as indicated were sub-cultured at 1:100 dilution and: A. grown for a further 3 hours at 37 °C, pelleted, re-suspended to an OD_600_ of 0.1 and 8 μL of sample was spotted onto LB agar or LB agar containing 2% SDS. Plates were incubated at 37 °C for 16 hours. or B. supplemented with 0.5% SDS (final concentration) and grown at 37 °C without shaking The optical density at 600nm was monitored every 20min using a plate reader. Error bars are ± SD, *n*=3 biological replicates.

**Figure S3.** Hydrophobic substitutions in the SufI signal peptide can rescue the transport defect of TatC^E103K^ and act in *cis* to suppress the SufI signal peptide R5K, R6K substitution. A. Overnight cultures of strain MC4100 Δ*amiA* Δ*amiC* Δ*tatABC* harboring pTAT101 producing TatABC^E103K^ along with pSUSufIss-mAmiA producing the indicated substitution in the SufI signal peptide were sub-cultured at 1:100 dilution and grown for a further 3 hours at 37 °C, pelleted, re-suspended to an OD_600_ of 0.1 and 8 μL of sample was spotted onto LB agar or LB agar containing 2% SDS. Plates were incubated at 37 °C for 16 hours. B. Overnight cultures of MC4100 Δ*amiA* Δ*amiC* Δ*tatABC* harboring pTAT101 (producing wild type TatABC) alongside either unsubstituted pSUSufIss-mAmiA (WT) or pSUSufIss-mAmiA encoding the indicated substitutions in the SufI signal peptide were sub-cultured at 1:100 dilution in the presence of 0.5% SDS (final concentration) and grown at 37 °C without shaking The optical density at 600nm was monitored every 20min using a plate reader.

**Figure S4.**
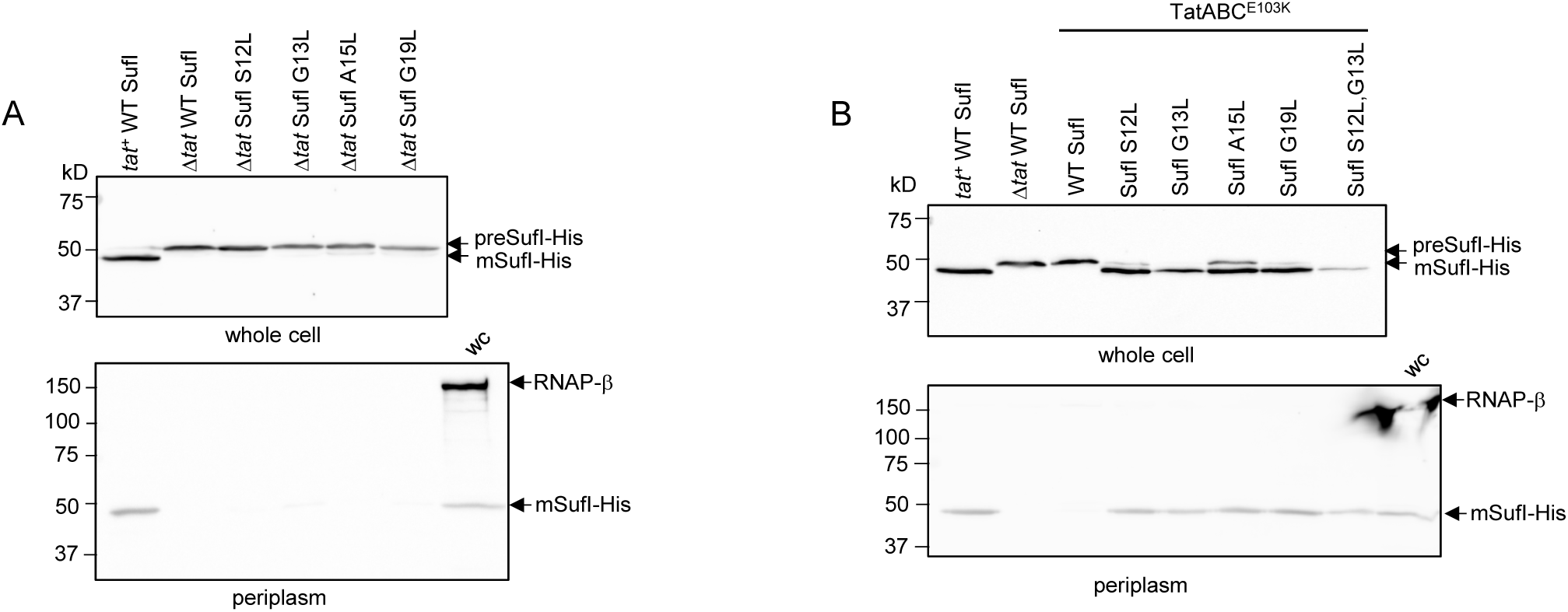
Analysis of SufI export mediated by signal peptide leucine substitutions in the TatC^E103K^ background and in a *tat* deletion strain. *E. coli* strain DADE co-producing his-tagged but otherwise native SufI or SufI with the indicated single leucine substitutions in the signal peptide (from a pQE80 plasmid) alongside either A. an empty plasmid vector (Δ*tat*) or B. wild type TatAB and TatC^E103K^ (from pTAT101) were grown to mid-log phase and fractionated into whole cell (upper panels) and periplasm (lower panels), then analyzed by Western blot with anti-6X His tag® or anti-RNA polymerase β subunit antibodies (cytoplasmic control protein. Strain DADE co-producing his-tagged but otherwise native SufI alongside either wild type TatABC (*tat*^+^ WT SufI) or an empty vector (Δ*tat* WT SufI) were used as positive and negative controls, respectively. wc – whole cell. Equivalent volumes of sample were loaded for each of the whole cell samples, and for each of the periplasmic samples.

**Figure S5.**
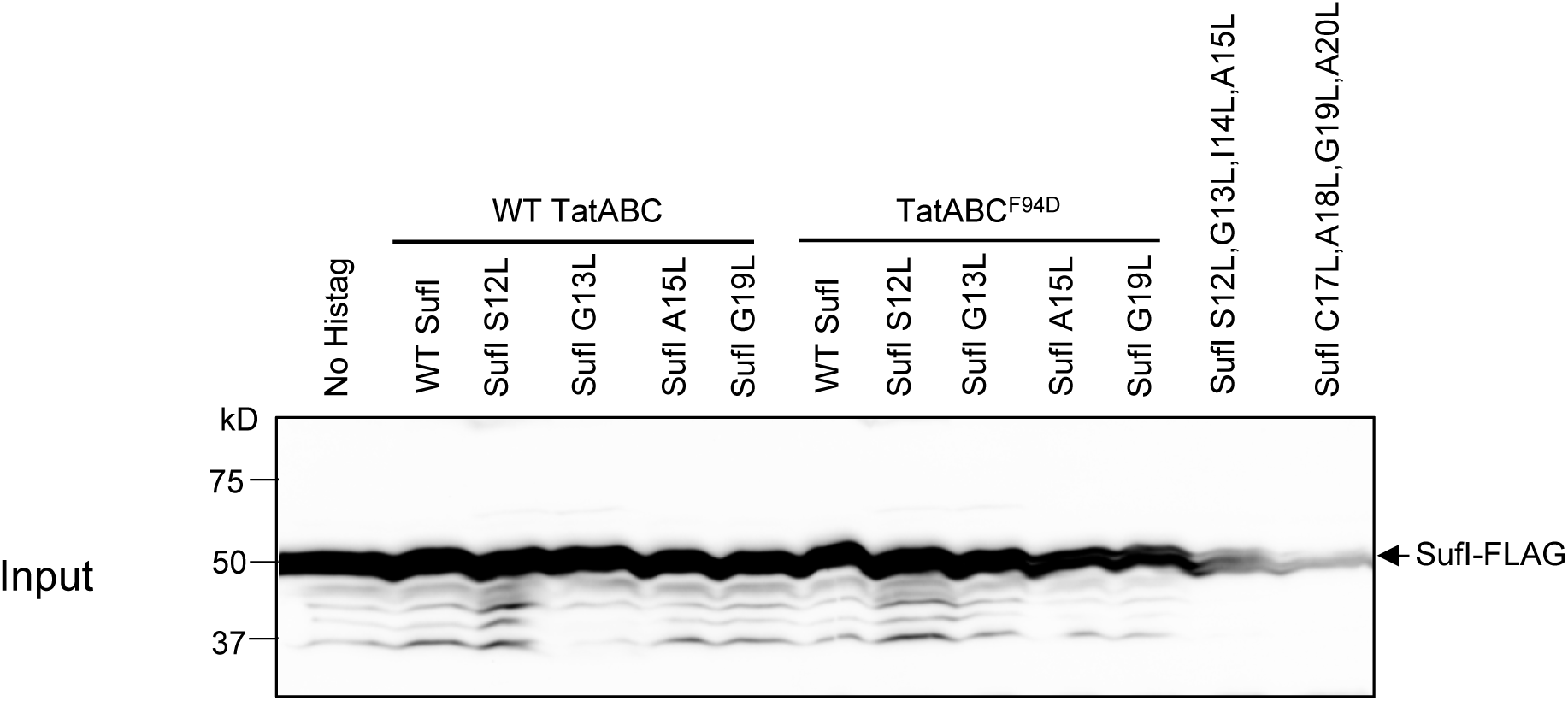
Detection of SufI-FLAG variants in input samples prior to Ni-affinity purification. Cells of strain DADE-P co-producing C-terminally FLAG-tagged SufI with its native signal peptide (WT SufI) or harboring the indicated leucine substitutions, alongside TatB and C-terminally His-tagged TatC or TatC^F94D^ were lysed and incubated with digitonin. 30 μL of the digitonin solubilised fraction was mixed with an equal volume of 2x Laemmli buffer and 10 μL of each sample was subject to SDS-PAGE (12% acrylamide) followed by western blot with anti-FLAG antibody.

